# Spaghettification: How extreme replichore imbalance impacts bacterial replication and morphology

**DOI:** 10.64898/2026.09.24.754080

**Authors:** Sha Cao, Filip Ilievski, Douglas L. Huseby, Magnus Johansson, Diarmaid Hughes, Gerrit Brandis

**Affiliations:** Department of Medical Biochemistry and Microbiology, Uppsala University, Uppsala, Sweden; Department of Cell and Molecular Biology, Uppsala University, Uppsala, Sweden; Uppsala Antibiotic Center, Uppsala University, Uppsala, Sweden

## Abstract

Bacterial chromosomes are typically organized into two similarly sized replichores to synchronize bidirectional replication and fusion within a Tus/*ter-*defined termination zone. Previous studies have suggested that replichore balance in both *Salmonella* and *E. coli* is maintained by selection for efficient and properly coordinated replication termination and chromosome segregation. However, there is a lack of understanding of the causal relationship between replichore asymmetry, cellular fitness, and replication-fork convergence, as well as how the termination machinery behaves when the two replichores differ substantially in length. Here we dissect the impact of replichore asymmetry by engineering 41 isogenic *Salmonella* inversion mutants spanning progressive imbalances, complemented by duplication-based asymmetries. Our constructed strains show that even cells with severely imbalanced chromosomes, where one replichore is 11-fold longer in size than the other (0.38 Mb vs. 4.48 Mb) are viable. However, relative fitness declined linearly with replichore size difference at approximately -0.17 per Mb. Additionally, highly imbalanced strains exhibited substantial division defects with severe cell elongation and lysis. Replication profiling showed that, even under extreme replichore asymmetry, forks consistently converge at the innermost *ter* sites of the shorter replichore and that two sequential Tus/*ter* barriers were sufficient for complete replication fork arrest. Together, these results establish a quantitative rule linking replichore asymmetry to fitness, define the termination position under imbalance, and reveal a dual Tus/*ter* barrier that safeguards termination when one replichore reaches the terminus region ahead of the other. Collectively these results connect chromosome architecture to replication mechanics and cell-division outcomes.

## Introduction

In most bacteria, chromosome replication initiates at a single origin (*oriC*) and proceeds bidirectionally until the two replication forks converge in the terminus region (1, 2). The two forks form independent replisomes that traverse opposite halves of the chromosome, generating two replichores. In the majority of species, these replichores are approximately equal in length (3), although notable exceptions occur in lineages that have undergone recent host adaptation, such as *Yersinia* species (4).

Replichore balance can be perturbed by large-scale chromosomal rearrangements, including duplications or insertions introduced into one replichore (*e.g.*, pathogenicity islands), or by asymmetrical inversions that displace the *oriC* or the terminus. Despite these potential disruptions, genomic surveys reveal that chromosomal inversions tend to occur symmetrically around *oriC*, and that isolates with balanced replichores exhibit a selective growth advantage (4, 5). Experimental work in *Escherichia coli* has further shown that when one replichore becomes less than half the length of the other, replication forks stall and DNA damage accumulates, underscoring the importance of replichore balance for robust growth and genome stability (6).

Replichore imbalance nevertheless arises frequently during chromosomal evolution, particularly through horizontal gene transfer, gene amplification, and inversion events associated with ecological or host adaptation (7–10). Previous studies examining the fitness effects of replichore imbalance have generally relied on individual engineered mutants or on strains representing discrete, non-systematic imbalance levels (11–14). These studies indicate that bacteria can tolerate modest replichore asymmetry without measurable fitness consequences. For example, *Salmonella enterica* strains carrying duplications resulting in one replichore being 0.22 Mb longer than the other (∼10%) displayed no significant growth defects (14). However, larger perturbations impose substantial fitness costs: an engineered *E. coli* strain with one replichore approximately three times as long as the other exhibited pronounced growth impairment, abnormal cell elongation, and defective division (15).

Despite these insights, a systematic and quantitative assessment of the relationship between replichore imbalance and bacterial fitness has been lacking. Here, we address this gap using *Salmonella enterica* serovar Typhimurium LT2 (*Salmonella* LT2), which possesses two nearly equal replichores of 2.45 Mb (left) and 2.40 Mb (right). We constructed a panel of strains spanning a continuous range of engineered replichore imbalance and quantified the resulting effects on bacterial fitness, replication dynamics, and cell proliferation. This systematic approach enables a comprehensive understanding of how bacterial chromosomes tolerate, or fail to tolerate, progressive deviations from replichore symmetry.

## Results

### Construction of inversion mutants with increasing degree of replichore imbalance

Isogenic *Salmonella* LT2 mutants with progressively imbalanced replichores were constructed using site-specific chromosomal inversions. As described in Materials and Methods, two truncated kanamycin-resistance gene fragments (*kan*), namely *kan\** (lacking the final 100 nucleotides) and \**kan* (lacking the promoter and the first 50 nucleotides), were inserted into the chromosome flanking the replication origin (*oriC*), with \**kan* and *kan*\* positioned in inverted orientation (Fig. 1A and B). Because these *kan* fragments are truncated, they do not confer kanamycin resistance, but they retain 645 bp of homologous sequence. Homologous recombination between the inverted \**kan* and *kan*\* sequences can result in a chromosomal inversion (16, 17), reconstituting an intact *kan* gene that confers kanamycin resistance (Fig. 1A and B). Inversion mutants were selected by plating overnight cultures of strains carrying both \**kan* and *kan*\* insertions on LA plates containing kanamycin. By positioning \**kan* and *kan*\* at defined chromosomal locations, we generated inversion mutants with predetermined degrees of replichore imbalance.

**Figure 1.**
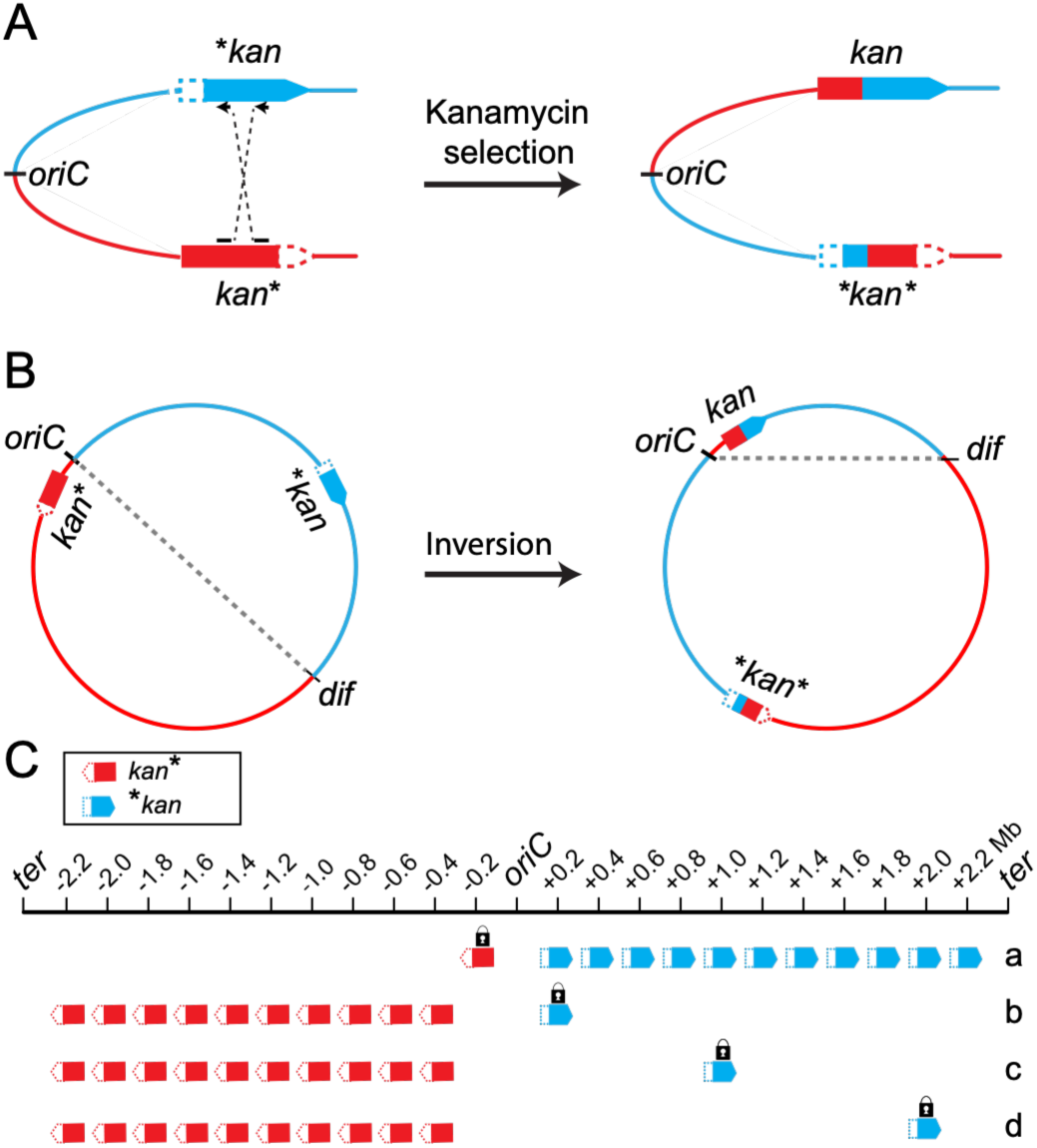
Overview of experimental design. (A) Schematic of the chromosomal inversion system. Two truncated *kan* fragments (*kan*\* and \**kan*) were inserted in inverted orientation flanking *oriC*. Homologous recombination between these inverted sequences can generate chromosomal inversion which reconstitutes an intact, functional *kan* gene. (B) Example of an imbalanced-replichore strain generated by inversion between *kan*\* at position -0.2 Mb and \**kan* at position +1.4 Mb. (C) Overview of inversion configurations. Four sets of strains were constructed to generate inversion mutants. In one set (a), *kan*\* was placed at a fixed chromosomal position; in the remaining three sets (b - d), \**kan* was placed at its respective fixed position. In all sets, the second *kan* fragment (*kan*\* or \**kan*) was inserted at one of ten additional positions. In total, 41 strains were constructed and used to select chromosomal inversion mutants.

Two sets of strains were constructed, comprising a total of 21 strains with \**kan* and *kan*\* inserted at different chromosomal locations. In the first set, *kan*\* was placed at a fixed position 0.2 Mb from *oriC* on the left replichore (−0.2 Mb), while \**kan* was inserted at 11 sites on the right replichore, spaced at 0.2 Mb intervals starting at position +0.2 Mb (Fig. 1C, set a). In the second set, \**kan* was fixed at +0.2 Mb on the right replichore, and *kan*\* was placed at 10 sites on the left replichore, also spaced at 0.2 Mb intervals starting from position -0.4 Mb since the combination with position -0.2 Mb was already constructed with the first set (Fig. 1C, set b). Chromosomal inversion mutants were successfully obtained for all 21 strains. Inversion frequencies ranged from 10^-6^ to 10^-9^ (SI Appendix, Table S1) (18), and one representative inversion mutant from each configuration was selected for further analysis.

The resulting mutants exhibit a progressive increase in replichore imbalance. Between successive mutants, the size of the shorter replichore decreases by approximately 0.2 Mb, while the longer replichore increases by the same amount (Fig. 2A and SI Appendix, Fig. S2). The extent of imbalance ranges from a mild inversion, such as INV(0Mb^-0.2/+0.2^), which maintains nearly balanced replichores (L/R: 2.38/2.48 Mb), to a highly imbalanced inversion in strain INV(−4Mb^-2.2/+0.2^) (L/R: 0.38/4.48 Mb), representing the most extreme configuration (Table 1). As a final quality control, long-read sequencing was performed for all 21 inversion mutants to confirm the intended inversions and exclude unintended secondary genome rearrangements. The successful isolation of all designated mutants demonstrates that all 21 strains, including those with the most extreme replichore imbalances, are viable.

**Figure 2.**
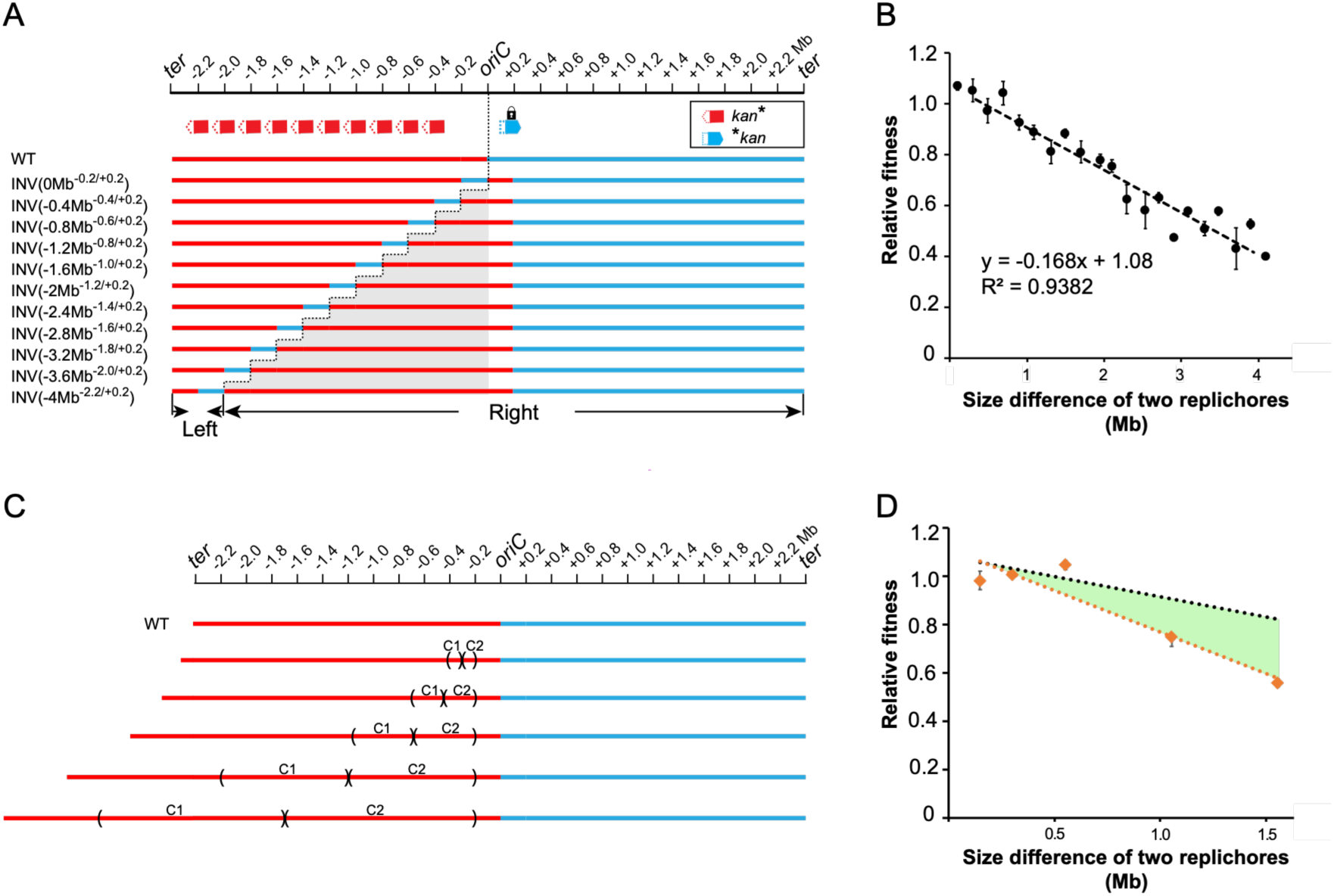
Relationship between replichore imbalance and bacterial fitness. (A) Schematic representation of replichore sizes following chromosomal inversion for one of the constructed strain sets. Inversions were generated between a fixed site at +0.2 Mb from *oriC* and one of eleven positions on the right replichore, spaced at 0.2 Mb intervals. (B) Correlation between replichore size asymmetry and bacterial fitness. The linear regression derived from inversion mutants was used to predict the expected fitness of chromosomal duplication mutants. (C) Schematic representation of replichore sizes following chromosomal duplications. (D) Correlation between replichore size asymmetry in duplication mutants and bacterial fitness. The black dashed line corresponds to the linear regression derived from inversion mutants from panel B. The green area indicates the additional fitness cost reduction in duplication mutants.

**Table 1.** Overview over replichore size differences and fitness effects in inversion strain with fixed insertions site at chromosomal positions -0.2 Mb and +0.2 Mb.

| Strain | Left replichore / Right replichore (Mb) | Replichore size difference (Mb) | Fitness ± S.D. |
| --- | --- | --- | --- |
| WT | 2.45 / 2.40 | 0.05 | 1.00 ± 0.02 |
| INV(0Mb<sup>-0.2/+0.2</sup>) | 2.38 / 2.48 | 0.10 | 1.10 ± 0.01 |
| INV(0.4Mb<sup>-0.2/+0.4</sup>) | 2.58 / 2.28 | 0.30 | 1.10 ± 0.04 |
| INV(+0.8Mb<sup>-0.2/+0.6</sup>) | 2.77 / 2.08 | 0.69 | 1.00 ± 0.05 |
| INV(+1.2Mb<sup>-0.2/+0.8</sup>) | 2.97 / 1.89 | 1.09 | 0.89 ± 0.03 |
| INV(+1.6Mb<sup>-0.2/+1.0</sup>) | 3.18 / 1.68 | 1.49 | 0.88 ± 0.02 |
| INV(+2Mb<sup>-0.2/+1.2</sup>) | 3.40 / 1.45 | 1.95 | 0.78 ± 0.02 |
| INV(+2.4Mb<sup>-0.2/+1.4</sup>) | 3.57 / 1.28 | 2.29 | 0.63 ± 0.06 |
| INV(+2.8Mb<sup>-0.2/+1.6</sup>) | 3.78 / 1.07 | 2.71 | 0.63 ± 0.02 |
| INV(+3.2Mb<sup>-0.2/+1.8</sup>) | 3.97 / 0.88 | 3.09 | 0.58 ± 0.01 |
| INV(+3.6Mb<sup>-0.2/+2.0</sup>) | 4.17 / 0.69 | 3.48 | 0.58 ± 0.02 |
| INV(+4Mb<sup>-0.2/+2.2</sup>) | 4.38 / 0.48 | 3.90 | 0.53 ± 0.02 |
| INV(-0.4Mb<sup>-0.4/+0.2</sup>) | 2.18 / 2.67 | 0.49 | 0.97 ± 0.04 |
| INV(-0.8Mb<sup>-0.6/+0.2</sup>) | 1.98 / 2.88 | 0.90 | 0.93 ± 0.03 |
| INV(-1.2Mb<sup>-0.8/+0.2</sup>) | 1.77 / 3.08 | 1.31 | 0.81 ± 0.05 |
| INV(-1.6Mb<sup>-1.0/+0.2</sup>) | 1.58 / 3.28 | 1.70 | 0.81 ± 0.04 |
| INV(-2Mb<sup>-1.2/+0.2</sup>) | 1.38 / 3.48 | 2.10 | 0.76 ± 0.02 |
| INV(-2.4Mb<sup>-1.4/+0.2</sup>) | 1.16 / 3.69 | 2.53 | 0.58 ± 0.07 |
| INV(-2.8Mb<sup>-1.6/+0.2</sup>) | 0.98 / 3.88 | 2.90 | 0.47 ± 0.01 |
| INV(-3.2Mb<sup>-1.8/+0.2</sup>) | 0.78 / 4.08 | 3.30 | 0.51 ± 0.03 |
| INV(-3.6Mb<sup>-2.0/+0.2</sup>) | 0.57 / 4.28 | 3.71 | 0.43 ± 0.08 |
| INV(-4Mb<sup>-2.2/+0.2</sup>) | 0.38 / 4.48 | 4.10 | 0.40 ± 0.03 |

### Cell fitness correlates with the degree of replichore size imbalance

We next examined whether replichore imbalance affects bacterial fitness and whether fitness defects scale with the degree of imbalance. Bacterial fitness was quantified for all inversion mutants based on changes in optical density at 600 nm (OD600) during exponential growth (Fig. 2B). No measurable fitness defect was detected in several mildly imbalanced strains, including INV(0Mb^-0.2/+0.2^), INV(+0.4Mb^-0.2/+0.4^), and INV(+0.8Mb^-0.2/+0.6^). In contrast, most inversion mutants exhibited fitness reductions of varying magnitude. Overall, fitness declined progressively as the size difference between the two replichores increased. The strain INV(−4Mb^-2.2/+0.2^), which carries the most extreme replichore imbalance, displayed the strongest growth defect, with an approximately 60% reduction in fitness relative to the wild-type strain. To quantitatively assess this relationship, bacterial fitness was plotted against the size difference between the two replichores for each inversion mutant. This analysis revealed a strong negative correlation between fitness and replichore size imbalance (R^2^ = 0.94). Analysis of the regression slope indicated a relative fitness cost of approximately 0.17 per megabase pair difference between the two replichores (Fig. 2B).

To test this relationship using an independent experimental strategy, an additional set of mutants with replichore imbalance was constructed by chromosomal duplication rather than inversion. As described in Materials and Methods, five isogenic strains were generated, each carrying a duplication of a DNA segment on the left replichore of approximately 0.1 Mb, 0.25 Mb, 0.5 Mb, 1 Mb, or 1.5 Mb (Fig. 2C). In these strains, the left replichore is extended by duplication, while the size of the right replichore remains unchanged. Fitness was measured experimentally using OD600-based growth analyses during exponential phase. No fitness defect was detected in strains carrying duplications of 0.1 Mb, 0.25 Mb, or 0.5 Mb, consistent with previous observations and indicating that small degrees of replichore imbalance are tolerated (14). In contrast, strains carrying larger duplications of 1 Mb and 1.5 Mb exhibited pronounced fitness defects. The duplication mutants showed greater fitness reductions than inversion mutants with corresponding replichore imbalances. Unlike inversion mutants, duplication mutants contain additional genetic material, and expression of genes encoded within the duplicated regions is likely to contribute to the observed fitness cost. In addition, increased gene dosage may perturb global regulatory networks and impose an additional metabolic burden, thereby further exacerbating the reduction in bacterial fitness (Fig. 2D).

### Replichore imbalance predicts fitness regardless of inversion location

Our previous analyses examined strains in which replichore imbalance was generated by chromosomal inversions with one inversion site fixed near *oriC* at either -0.2 Mb or +0.2 Mb. To determine whether the relationship between fitness and replichore imbalance depends on the specific chromosomal locations involved in the inversion, we constructed two additional sets of isogenic inversion mutants. In these sets, one inversion site was fixed on the right replichore at either +1.0 Mb or +2.0 Mb, while the second site was varied across ten positions on the left replichore, ranging from -0.4 Mb to -2.2 Mb in 0.2 Mb intervals (SI Appendix, Fig. S2A and B). These mutants were generated using the same approach described above: insertion of kan* and *kan at designated chromosomal sites followed by selection of inversion mutants on kanamycin plates (Fig. 1C, sets c and d).

As in the previous sets, the degree of replichore imbalance in these strains increased progressively across the constructed series. Relative fitness for each strain was quantified based on OD600 measurements during exponential growth. Most mutants with small degrees of replichore imbalance exhibited no or only minor fitness defects. Consistent with earlier observations, increasing replichore imbalance resulted in progressively larger fitness defects (SI Appendix, Table S2). The correlation between fitness and the size difference of the two replichores remained evident: strains with greater replichore size asymmetry exhibited more pronounced fitness losses. When the fitness values of all 41 inversion mutants were plotted against the size difference between their two replichores, the relationship showed a strong overall negative correlation (R^2^ = 0.82) with a slope of -0.14 per Mb (SI Appendix, Fig. S2C). These results indicate that replichore size asymmetry predicts reduction in cellular fitness regardless of inversion location.

### Replication terminates at the innermost *ter* site of the shorter replichore

In bacteria, DNA replication terminates when the two replication forks meet within the terminus region, which lies opposite *oriC* and is flanked by the innermost replication termination sites (*ter* sites). In strains with balanced replichores, the replication forks typically reach this region at approximately the same time (19). Enterobacteriaceae such as *E. coli* and *Salmonella* contain multiple *ter* sites distributed across the genome (1, 20), and when bound by the Tus protein, these sites arrest replication forks in a polar manner (21, 22). If one replichore completes replication earlier than the other, the Tus/*ter* complex can trap the leading replication fork, preventing it from progressing into the opposite replichore. This raises a fundamental question for strains with strongly imbalanced replichores, particularly those with extreme asymmetry: Can the replication fork of the shorter replichore be reliably halted at its innermost *ter* site and remain arrested until the longer replichore completes replication? If arrest is incomplete or bypass occurs frequently the effective lengths of the replichores would differ from their nominal sizes. In *Salmonella*, ten *ter* sites are distributed across the chromosome, with the innermost *ter* sites being *ter9* and *ter1* for the left and right replichores, respectively (Fig. 3A) (23).

**Figure 3.**
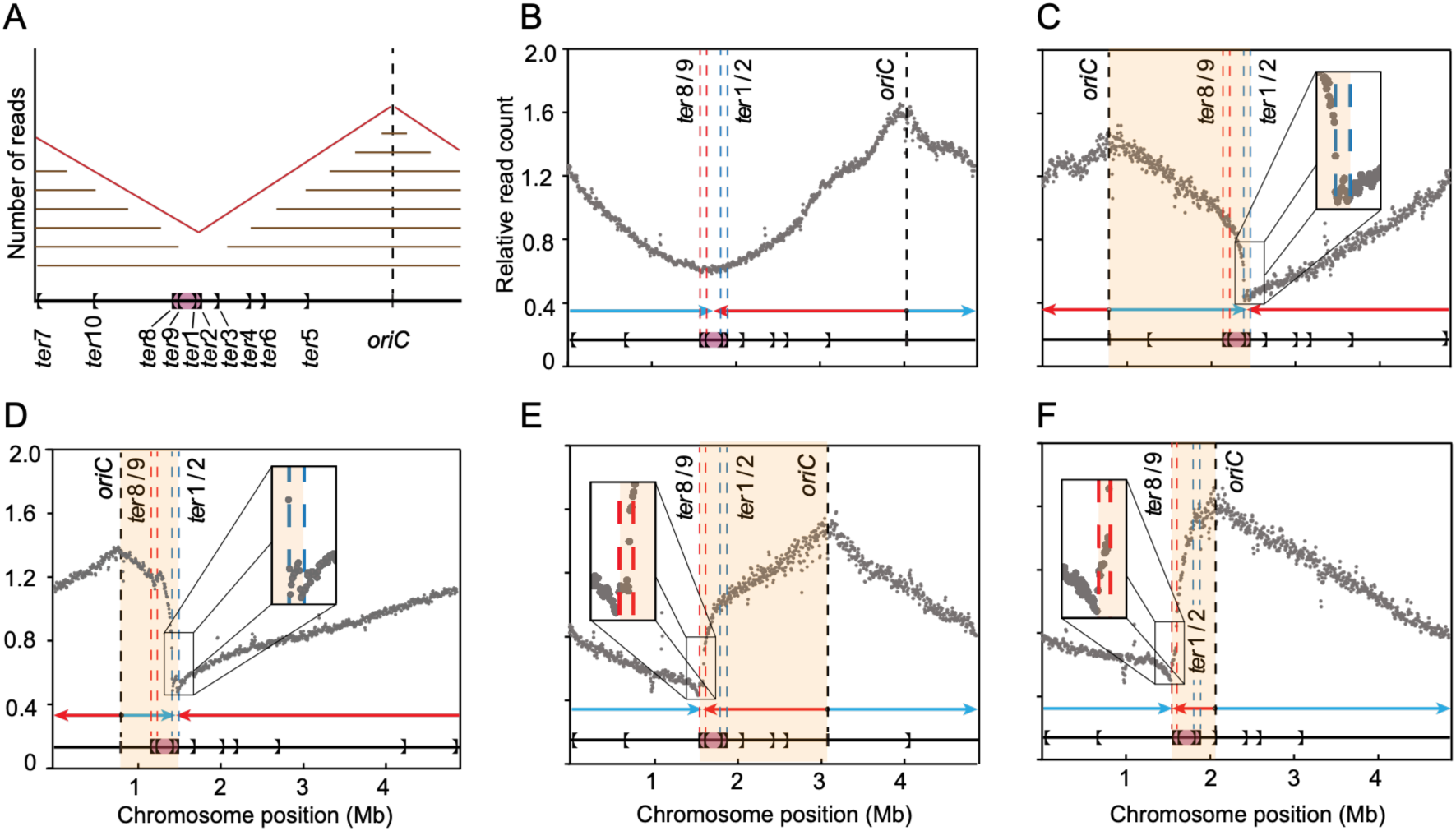
Replication profile analysis of inversion mutants. (A) Schematic map of the *Salmonella* LT2 chromosome showing the ten *ter* sites. The purple-shaded region indicates the termination buffer zone between the *ter* sites pairs that effectively block all replication forks (*ter8/9* and *ter1/2*). (B - F) Marker frequency analysis of representative strains. Relative read count (grey dots) from exponentially growing cells, normalized to stationary-phase cells, as a function of chromosome position. Replication forks originating from the two replichores are indicated by red and cyan arrows. The position of *oriC* is marked by a black dashed line. The innermost *ter* sites for each replichore are marked with dashed lines in the same colour as the corresponding replication fork. The region corresponding to the shorter replichore is shaded in yellow. (B) Wild-type LT2, (C) INV(+2Mb^-0.2/+1.2^), (D) INV(+4Mb^-0.2/+2.2^), (E) INV(−2Mb^-1.2/+0.2^), and (F) INV(−4Mb^-2.2/+0.2^).

To address this question, we selected five representative strains for replication profiling: INV(+2Mb^-0.2/+1.2^), INV(+4Mb^-0.2/+2.2^), INV(−2Mb^-1.2/+0.2^), INV(−4Mb^-2.2/+0.2^), and the wild type (WT). Replication profiles were obtained using marker frequency analysis (MFA), which relies on deep sequencing of short reads from exponentially growing cultures. In the WT strain, MFA revealed the expected symmetric V-shaped profile, with maximal read coverage at *oriC* and a distinct minimum between *ter9* and *ter1*, confirming convergence of the two forks at the terminus region (Fig. 3B). In all inversion mutants examined, the replication profiles displayed a clear minimum at the innermost *ter* site of the shorter replichore. In strains INV(+2Mb^-0.2/+1.2^) and INV(+4Mb^-0.2/+2.2^), the minima localized to the *ter1/2* region (Fig. 3C and D), whereas in strains INV(−2Mb^-1.2/+0.2^) and INV(−4Mb^-2.2/+0.2^), minima were positioned at *ter8/9* (Fig. 3E and F). These results demonstrate that even in highly imbalanced chromosomes, replication forks consistently converge at the innermost *ter* site of the shorter replichore. Notably, two distinct minima were observed in strains INV(+2Mb^-0.2/+1.2^), INV(+4Mb^-0.2/+2.2^), and INV(−2Mb^-1.2/+0.2^). This pattern suggests that while replication forks are efficiently halted at the innermost *ter* site, a subset of forks may bypass the initial arrest and proceed until they encounter the second innermost *ter* site (Fig. 3C-E).

### Complete replication fork arrest requires two Tus/ter barriers

In wild-type bacteria with balanced replichores, replication terminates between the two innermost *ter* sites *ter9* and *ter1*. Each of these sites is in close proximity to a secondary *ter* site forming pairs: *ter8/9* on one side and *ter1/2* on the other (Fig. 3B). We asked whether both members of each *ter*-site pair are required to ensure complete replication fork arrest. To address this, we deleted either the innermost (*ter9*, *ter1*) or the second innermost (*ter8*, *ter2*) *ter* sites in four strains used for the replication termination analyses, INV(−4Mb^-2.2/+0.2^), INV(−2Mb^-1.2/+0.2^), INV(+2Mb^-0.2/+1.2^), and INV(+4Mb^-0.2/+2.2^) (Fig. 4A). However, two cases could not be examined for technical reasons: (i) deletion of *ter8* in strain INV(−2Mb^-1.2/+0.2^), because *ter8* also regulates expression of *tus*, and its removal would compromise Tus-dependent fork arrest at all *ter* sites; and (ii) deletion of *ter* sites in strain INV(−4Mb^-2.2/+0.2^), which consistently resulted in large-scale chromosomal rearrangements, likely because loss of these termination barriers in a chromosome with extreme replichore imbalance generates lethal replication-segregation conflicts.

**Figure 4.**
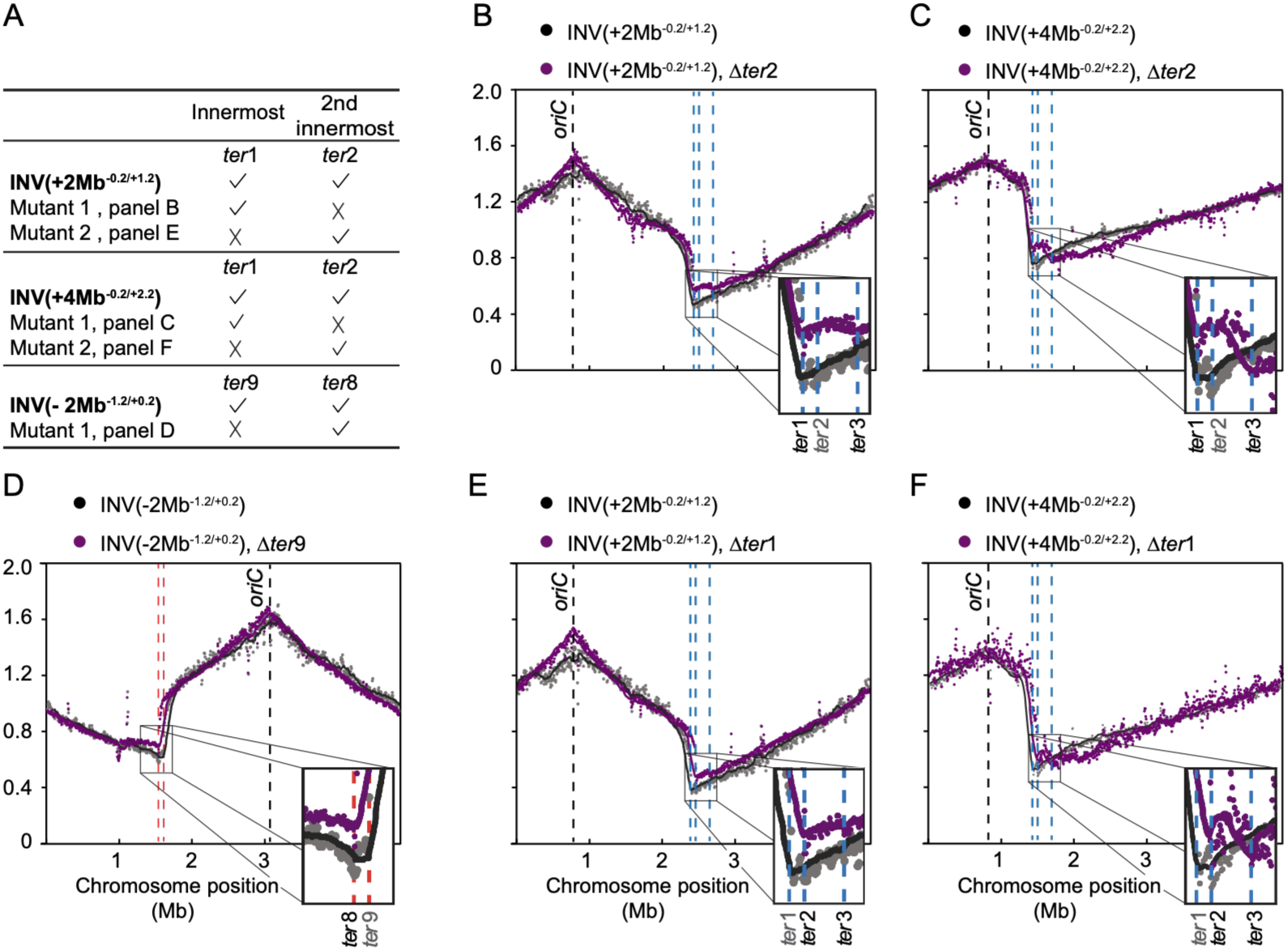
Replication profiles after deletion of the innermost *ter* sites of the shorter replichore. (A) List of parent strains and their corresponding *ter*-deletion mutants. (B - F) Marker frequency analysis of parent strains (grey dots) and their *ter*-deletion derivatives (purple dots). Relative read counts from exponentially growing cells, normalized to stationary-phase cells, are plotted against chromosome position. Trend lines are shown as solid curves (parent: black; mutant: purple). The position of *oriC* is indicated by a black dashed line. Relevant *ter* sites of the shorter replichore are marked with dashed lines; deleted *ter* sites in the mutant profiles are labelled in light grey font. (B) INV(+2Mb^-0.2/+1.2^): parent and Δ*ter2*, (C) INV(+4Mb^-0.2/+2.2^): parent and Δ*ter2*, (D) INV(−2Mb^-1.2/+0.2^): parent and Δ*ter9* (E) INV(+2Mb^-0.2/+1.2^): parent and Δ*ter1*, and (F) INV(+4Mb^-0.2/+2.2^): parent and Δ*ter1*.

Deletion of the innermost *ter* sites (*ter9* in strain INV(−2Mb^-1.2/+0.2^), and *ter1* in strains INV(+2Mb^-0.2/+1.2^) and INV(+4Mb^-0.2/+2.2^)) shifted the primary minimum to the second innermost *ter* site (Fig. 4D - F). This demonstrates that the majority of approaching replication forks are efficiently halted at the first *ter* site they encounter, regardless of which specific *ter* site occupies that position. Despite these shifts, MFA profiles for these mutants generally displayed two distinct minima, typically positioned at the second innermost and third innermost *ter* sites (e.g., *ter2* and *ter3*; Fig. 4E and F). Consistent with these results, MFA revealed that deletion of the second innermost *ter* site (*ter2*) in strains INV(+2Mb^-0.2/+1.2^) and INV(+4Mb^-0.2/+2.2^) did not alter the position of the primary minimum in the replication profiles. However, the secondary minimum shifted from *ter2* to *ter3* (Fig. 4B and C). This pattern reinforces the conclusion that complete replication termination frequently requires two Tus/*ter*-mediated arrests, with the first barrier halting the fork and the second ensuring full convergence and termination.

### Severe cell-division defects arise in cells with imbalanced replichores

Cell proliferation is a tightly coordinated process involving chromosome replication, nucleoid segregation, cell elongation, and division (24–27). Replichore imbalance alters the timing and progression of replication forks, and previous work has shown that *E. coli* cells with asymmetric replichores exhibit elongation and inhibited division (15). To determine whether similar defects occur in our *Salmonella* inversion mutants, we examined four strains - WT, INV(−1.2Mb^-0.8/+0.2^), INV(−2Mb^-1.2/+0.2^), and INV(−4Mb^-2.2/+0.2^) - using time-lapse microscopy.

A prominent feature across all mutants was cell filamentation, which increased in severity with greater replichore imbalance. WT cells displayed uniformly sized, ellipsoidal morphology (Fig. 5A, Video S1A). In contrast, strain INV(−1.2Mb^-0.8/+0.2^), which carries only mild replichore asymmetry, exhibited branched cells with multiple polar ends and modest filamentation (Fig. 5B, Video S1B). In strains INV(−2Mb^-1.2/+0.2^) and INV(−4Mb^-2.2/+0.2^), the majority of cells became severely elongated, and a substantial fraction ultimately lysed. A small population of rounded, nongrowing cells was also observed, potentially representing chromosome-free cells due to asymmetric cell division (Fig. 5C and D, Video S1C and D). Cells of strain INV(−4Mb^-2.2/+0.2^), which carries the most extreme imbalance, formed highly elongated, spaghetti-like filaments that accumulated along the edges of the microfluidic chamber and substantial cell lysis was observed (Fig. 5D and Video S1D). A live/dead assay further confirmed the presence of elongated dead cells within the population of non-lysed cells in the INV(−4Mb^-2.2/+0.2^) strain (Fig. 5E). Collectively, these observations indicate that cell division is severely disrupted in cells with markedly imbalanced replichores.

**Figure 5.**
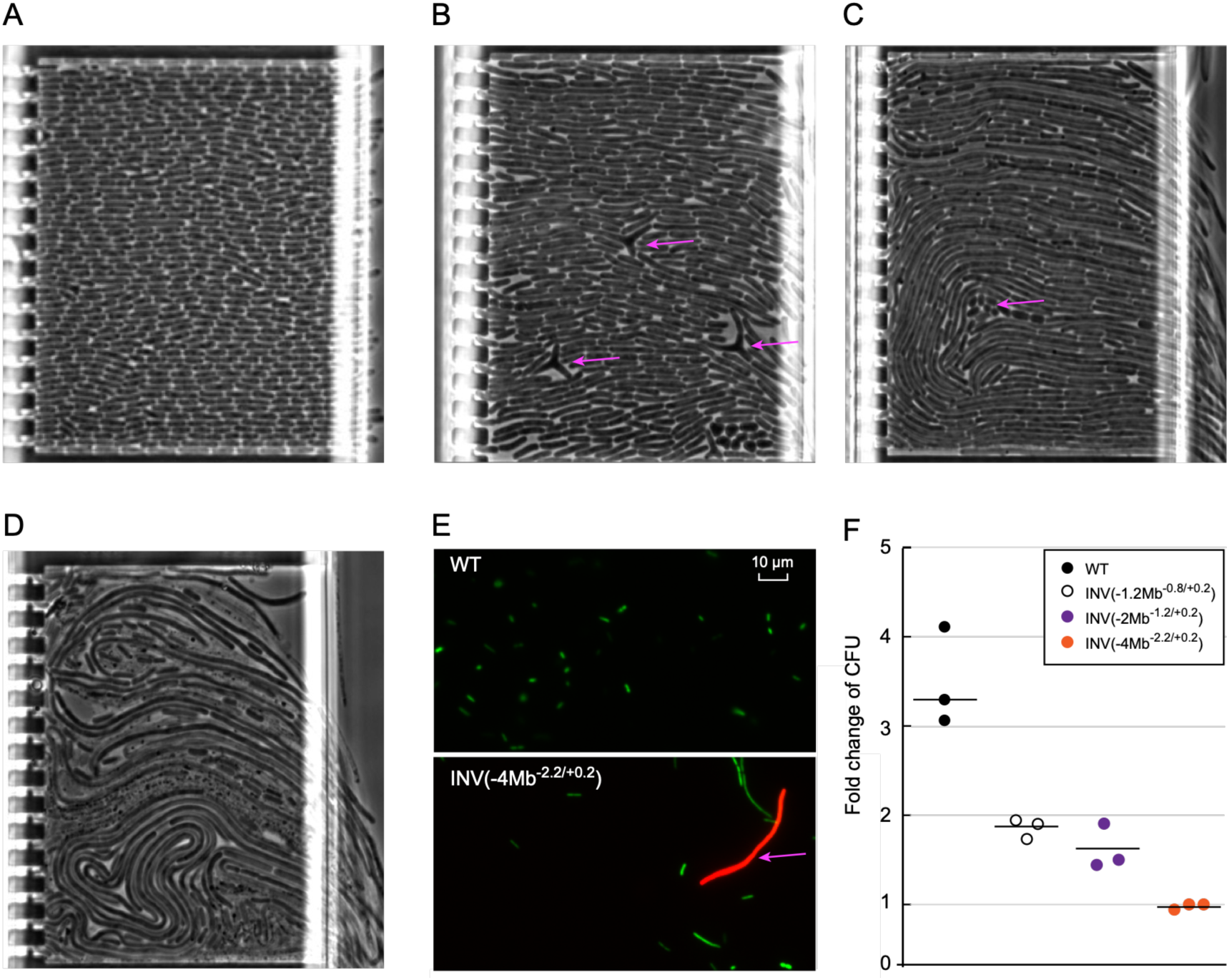
Cell morphology and viability of representative inversion mutants. (A - D) Time-lapse microscopy images illustrating morphological changes in strains with increasing replichore imbalance. (A) Wild-type cells display a uniform, ovoid morphology. (B) INV(−1.2Mb^-0.8/+0.2^) cells show moderate elongation and occasional branched structures (purple arrows). (C) INV(−2Mb^-1.2/+0.2^) cells exhibit two abnormal subpopulations: small, rounded cells that fail to elongate (purple arrows) and elongated filamentous cells. (D) INV(−4Mb^-2.2/+0.2^) cells display extensive filamentation (“spaghettification”) and widespread lysis, visible as grey flocculent material containing black granular debris. (E) Live/dead (green/red) fluorescence imaging of wild type (top) and INV(−4Mb^-2.2/+0.2^) (bottom). (F) Fold change in colony-forming units (CFU) between OD600 = 0.04 and 0.08.

To quantify these defects, we measured the change in colony-forming units (CFU) during early exponential growth. Between OD600 values of 0.04 and 0.08, CFU in the WT increased approximately three-fold. In contrast, only modest increases of 1.6-fold and 1.3-fold were observed in the INV(−1.2Mb^-0.8/+0.2^) and INV(−2Mb^-1.2/+0.2^) strains, respectively, and no increase in CFU was detected in strain INV(−4Mb^-2.2/+0.2^) (Fig. 5F). These results strongly support the conclusion that bacterial cells with imbalanced replichores fail to complete normal cell division during growth.

## Discussion

Circular bacterial chromosomes are typically organized into two replichores, with bidirectional replication initiating at a single origin (*oriC*) and proceeding until the forks fuse in a terminus region. In many bacterial species, such as *Salmonella*, polar Tus/*ter* barriers form a replication trap that prevents the replication fork from progressing beyond this region (20). This symmetrical architecture is thought to minimize the time to complete DNA replication and thereby support optimal growth. Previous work demonstrated the importance of replichore balance on bacterial fitness and cell proliferation (14, 15, 28, 29) and linked genome rearrangements to lifestyle in specific *S. enterica* serovars (8). Our study advances this area by employing a carefully designed, experimentally controlled approach: we engineered a large, stepwise panel of inversions (and an auxiliary set of duplications) to define replichore lengths *a priori* while holding gene content constant in the inversion series, isolating the effects of replichore asymmetry on fitness, replication termination, and cell division.

We generated and verified 41 mutants spanning a wide range of replichore imbalances, including strains with extreme asymmetry (Table 1 and SI Appendix, Table S2). The shorter replichore of the most imbalanced strain is only 0.38 Mb long which is 11-fold shorter than the longer replichores (4.48 Mb). Although strains with pronounced replichore imbalances displayed significant fitness defects, they remained viable. Mild asymmetries yielded little or no detectable defect, consistent with earlier observations that limited imbalance can be tolerated without a clear fitness cost (14). Across the inversion series, relative fitness declined linearly with replichore size difference (slope - 0.14 per Mb), establishing a quantitative rule that connects chromosome architecture to population-level growth (Fig. 2A and B). It is important to note, however, that these fitness estimates are derived from exponential-phase optical density measurements, which report biomass accumulation. Subsequent experiments showed that in strains with extreme replichore imbalance, optical density rise does not accurately reflect the production of viable daughter cells indicating that the fitness costs inferred from growth curves likely underestimate the true severity of the defect at the highest levels of asymmetry (Fig. 5F). Our experiments with duplication mutants showed that duplication-based asymmetries produced more severe fitness loss than inversions. We speculate that this is due to an added burden from changes in the gene dosage and global regulatory disruption (Fig. 2C and D). Together, these data indicate that replichore imbalance imposes an inherent replication-schedule cost, and that additional DNA exacerbates the cost by perturbing gene dosage and regulon balance.

At the level of replication mechanics, marker frequency analysis revealed a consistent and informative pattern: even under the most extreme asymmetry (0.38 Mb vs. 4.48 Mb), forks converged at the innermost *ter* of the shorter replichores (Fig 3) in accordance to the replication-fork trap model (30). Deletion analyses then uncovered a second, key principle: two sequential Tus/*ter* barriers are sufficient for full fork arrest. Removing the innermost site shifted the primary MFA minimum to the second-innermost site, while removing the second-innermost site displaced the secondary minimum to the third-innermost site accordingly, indicating that a first barrier halts most forks, whereas a second barrier ensures complete convergence and prevents escape into the opposite replichores (Fig. 4). We did not observe escape beyond the second barrier, suggesting either (i) robust containment by the dual trap or (ii) inviability of cells in which forks progress beyond the second site (*e.g.,* due to failure of decatenation or fatal interference with late cell-cycle processes) (1, 31). The observed two-minimum profiles are consistent with occasional bypass of the first site followed by capture at the second, reinforcing the concept of a layered fail-safe within the terminus region (Fig. 4C and F) (30). These results are in accordance to a previous study where two *ter* sites were sufficient to halt replication fork progression in more modestly imbalanced chromosomes (31). The terminus region itself appears to buffer a certain degree of asymmetry. In *Salmonella* LT2, the span between the second innermost *ter* sites of each replichore (*ter2* and *ter8*) is ∼0.32 Mb, providing a zone that tolerates limited over-travel of the fork from the shorter arm. This buffer model explains why minor imbalances do not necessarily impair fitness, and it hints at an evolutionary rationale for multiple *ter* sites: by providing redundant, polar barriers, the system maintains termination near the chromosome’s division site even when replication schedules are perturbed.

Replichore imbalance also disrupts post-replicative and division processes, consistent with earlier observations in *E. coli* (15). When one arm requires substantially longer to replicate than the other, the resulting asynchrony interferes with the coordination of decatenation, nucleoid segregation, FtsZ-ring assembly, and septum formation, leading to filamentation and lysis in the most imbalanced strains (24–27, 32). Filamentation may be aggravated if the altered *oriC*/*ter* geometry brings SlmA binding sites into inhibitory proximity to the division plane, further impeding productive Z-ring formation (33, 34). Our CFU measurements during exponential phase confirm that the most imbalanced strain fails to increase viable counts despite an OD600 rise, linking replication-schedule disruption to division failure and death (Fig. 5F)

Collectively, these findings connect genome architecture to replication mechanics and cell physiology. First, the linear fitness-imbalance rule (−0.17 per Mb) quantifies the long-standing intuition that symmetric replichores are beneficial. Second, termination is fixed by the shorter replichore’s innermost *ter*, revealing how the fork-trap geometry resolves severe schedule asymmetries. Third, a dual Tus/*ter* barrier is sufficient to maintain the stability of termination when one arm substantially outpaces the other. Forth, the ∼0.32 Mb region between the second innermost *ter* sites of each replichore acts as a buffer when replication schedules are perturbed. Fifth, severe replichore imbalances disrupts post-replicative and division processes leading to division failure and cell lysis. Finally, by integrating inversion-based (no added DNA) with duplication-based (added DNA) imbalances, we separate the geometric costs of asymmetry from the dosage/regulatory costs that accompany additional DNA.

In summary, balanced replichores emerge as a product of evolutionary optimization that synchronizes fork schedules, stabilizes termination, and enables robust division. Mild deviations can be buffered by the Tus/*ter* fork-trap, but substantial asymmetry pushes the system into a regime characterized by prolonged stalling at the first barrier, reliance on a second barrier for complete fork arrest, and, ultimately, a breakdown in cell-division coordination that leads to severe cell elongation and death.

## Materials and Methods

### Growth conditions

All media and chemicals were purchased from Merck unless otherwise specified. Bacteria were routinely cultured in LB medium (1% tryptone, 0.5% yeast extract [Oxoid, England] and 1% NaCl) or on LA plates (LB with 1.5% bacteriological agar, Oxoid) supplemented with appropriate antibiotics and incubated at 37°C.

### Strain constructions

All strains used in this study are derivatives of *Salmonella enterica* serovar Typhimurium strain LT2 (35). Two kanamycin cassettes with different truncations, \**kan* and *kan*\*, were linked to *bla* (*bla*-\**kan*) or *cat* (*cat*-*kan*\*), respectively. The *bla* encodes a β-lactamase, providing resistance to ampicillin, while the *cat* gene encodes a chloramphenicol acetyltransferase, conferring chloramphenicol resistance. These markers enable the selection of strains that have been successfully transformed with either cassette (SI Appendix, Fig. 3A). The insertions of *bla*-\**kan* and *cat*-*kan*\* on the chromosome were performed though lambda red recombineering (36). Convergent intergenic regions were chosen for all insertion sites to minimize the impact on native gene expression. Following the *bla*-\**kan* and *cat*-*kan*\* insertions, three independent cultures of each constructed strain were plated on LA plates containing kanamycin to select for chromosome inversion mutants. Colony counts were recorded and used to calculate the inversion frequency. PCR amplifications of the inversion junctions were performed to verify successful generation of inversions. One strain from each inversion event was selected for further analysis. The selected strains were whole-genome sequenced both by short-read and long-read sequencing to confirm that no unintended mutations or chromosome rearrangements were selected during construction process.

Chromosomal duplications were generated by transforming the strain LT2 Δ*dap*A with a PCR amplified *dap*A fragment flanked by 45 nucleotide homology arms corresponding to the ends of the chromosomal region targeted for duplication. Strain LT2 Δ*dap*A is auxotrophic to diaminopimelic acid (DAP) and therefore cannot grow on LA plates. Thus, the chromosomal duplication created by the *dapA* insertion is maintained while strain is grown in LB medium or on LA plates (SI Appendix, Fig. S3B) (37). All strains constructed in this study are listed in SI Appendix, Table S3.

### PCR

PCRs for recombineering were done using Phusion high-fidelity PCR master mix (New England Biolabs), according to the manufacturer’s protocol. Taq polymerase-based master mix (Thermo Fischer Scientific) were used for verifying inversions and preparing DNA for local sequencing. Oligonucleotides for PCR were synthesized at MerckReactions were run on a model S1000 thermal cycler from Bio-Rad. All recombineering and strain verification primers are listed in SI Appendix, Table S4.

### Whole genome sequencing

Bacterial genomic DNA was prepared by using MasterPure DNA & RNA purification kit (Epicentre). The concentration of the extracted DNA was assessed using the Qubit 2.0 fluorometer (Invitrogen). The short reads sequencing was performed on site using Miseq (Illumina Inc.) while the long reads sequencing was carried on MinION Mk1C (Oxford Nanopore Technologies). The sequencing data was aligned and analyzed using the software CLC Genomics Workbench V11 (Qiagen) and Unicycler (38).

### Growth measurements

Doubling time of strains were measured by monitoring the increase of optical density with a Bioscreen C machine (Oy Growth curves AB Ltd). Fresh bacterial cells were prepared in 0.9% NaCl by scraping from LA plates and applied directly to honeycomb microtiter plates (Oy Growth curves AB Ltd) supplemented with LB medium. The honeycomb plates were then placed in a Bioscreen machine and grown at 37 °C for 18 hours with continuous shaking. The optical density of cells was measured every 5 minutes at OD600nm. The doubling time of bacteria were calculated by plotting the OD600 value against the growth time at exponential growth phase (OD600nm = 0.04 to 0.08). A trendline was applied and the slope of the trendline was used to calculate the doubling time. The equation used was: doubling time = ln(2)/slope (minutes).

### Marker frequency analysis (MFA)

Exponentially growing bacteria were prepared in liquid cultures (OD600nm ≈ 0.3). Genomic DNA was purified and sent to BGI (BGI Genomics Co., Ltd) for short read deep sequencing. Sequence mapping and read depth analysis were performed using CLC genomic workbench, and the read depth in 10 kb windows with 5 kb overlap was calculated using Perl. To account for differences in sequence depth across the chromosome that are due to sequencing biases, the read depth of exponentially growing bacteria was normalized to that of the same strain in stationary phase. After normalization, the read coverage was used to represent the replication dynamics of chromosome (15, 17).

### Time-lapse microscopy

Microfluidic chip design and manufacturing is described elsewhere (39). The microfluidic chip consists of a molded polydimethylsiloxane (PDMS; Sylgard 184) elastomer layer covalently bonded to a borosilicate cover glass (no. 1.5 thickness, 22 × 40 mm, VWR). Tubing (VWR, TYGON VERNAAD04103) was attached to the chip using metal tubing connectors. Media flow was controlled using an in-house made Arduino-based pressure regulator (Elveflow).

For the experiments, bacterial cells were scraped from LA plates and suspended in 0.9% NaCl to a density of OD600 ≈ 1. Then 20 µl of the cell suspension was inoculated into 2 ml of pre-warmed LB medium and allowed to grow to exponential growth phase (1.5 hours for WT, 2-3 hours for mutants). Cells in the exponential growth phase were subsequently used for the single cell analysis. 15 mL Falcon tubes served as media and culture reservoirs, positioned level with the microscopy stage and connected to the chip via Elveflow adapters and tubing (VWR, TYGON VERNAAD04103). Metal connectors (23-gauge, 14 mm, 90°-bent; New England Small Tubing) were used at the tubing-to-port junctions. Prior to time-lapse imaging, chips were wetted with LB media supplemented with pluronics. Following wetting, cells were loaded at 200 mbar, with a counter-pressure of 50 mbar at media-in ports. After cell loading, all pressures were decreased and growth media was supplied at 150 mbar.

Phase contrast timelapse microscopy was performed every 2 min over 4 hours using an inverted Nikon Ti2-E microscope fitted with a CFI Plan Apo Lambda 100×/1.45 objective. The setup was housed within an OKOlab environmental chamber (model H201-ENCLOSURE) paired with a temperature control unit (H201-TUNIT-BL) to keep the sample at a stable 37 ± 2 °C throughout experiments. Phase contrast time-lapse images were acquired with a Hamamatsu Orca Quest camera, coupled to the microscope through a 0.7× demagnifying camera adapter.

15 mL Falcon tubes served as macrofluidic reservoirs, positioned level with the microscopy stage and connected to the chip via Elveflow adapters and tubing (VWR, TYGON VERNAAD04103). Metal connectors (23-gauge, 14 mm, 90°-bent; New England Small Tubing) were used at the tubing-to-port junctions. Prior to experiments, chips were wetted with growth media in the following port sequence: ports 5.1, 5.2, and 6.0 at 800 mbar, port 2.0 at 800 mbar, and ports 2.1 and 2.2 at 700 mbar, with flow regulated by the OB1-Mk3 pressure controller (Elveflow). Following wetting, pressure to ports 5.1, 5.2, and 6.0 was discontinued. Bacterial cells were loaded via port 2.0 at 120 mbar, with a counter-pressure of 50 mbar applied at ports 2.1 and 2.2. Loading pressure could be incrementally raised as needed to ensure complete microchamber filling. During experiments, media were supplied through ports 2.1 and 2.2 at 80 – 100 mbar, with no pressure applied to any other ports.

### Fluorescence microscopy

Strains were grown to exponential growth phase and then stained using the LIVE/DEAD BacLight Bacterial Viability Kit (Invitrogen, ThermoFisher Scientific) according to the manufacturer’s instructions. Fluorescence imaging was performed using a Nikon Eclipse 90i microscope, and cell images were acquired accordingly.

## Supplementary Information

**Fig. S1.**
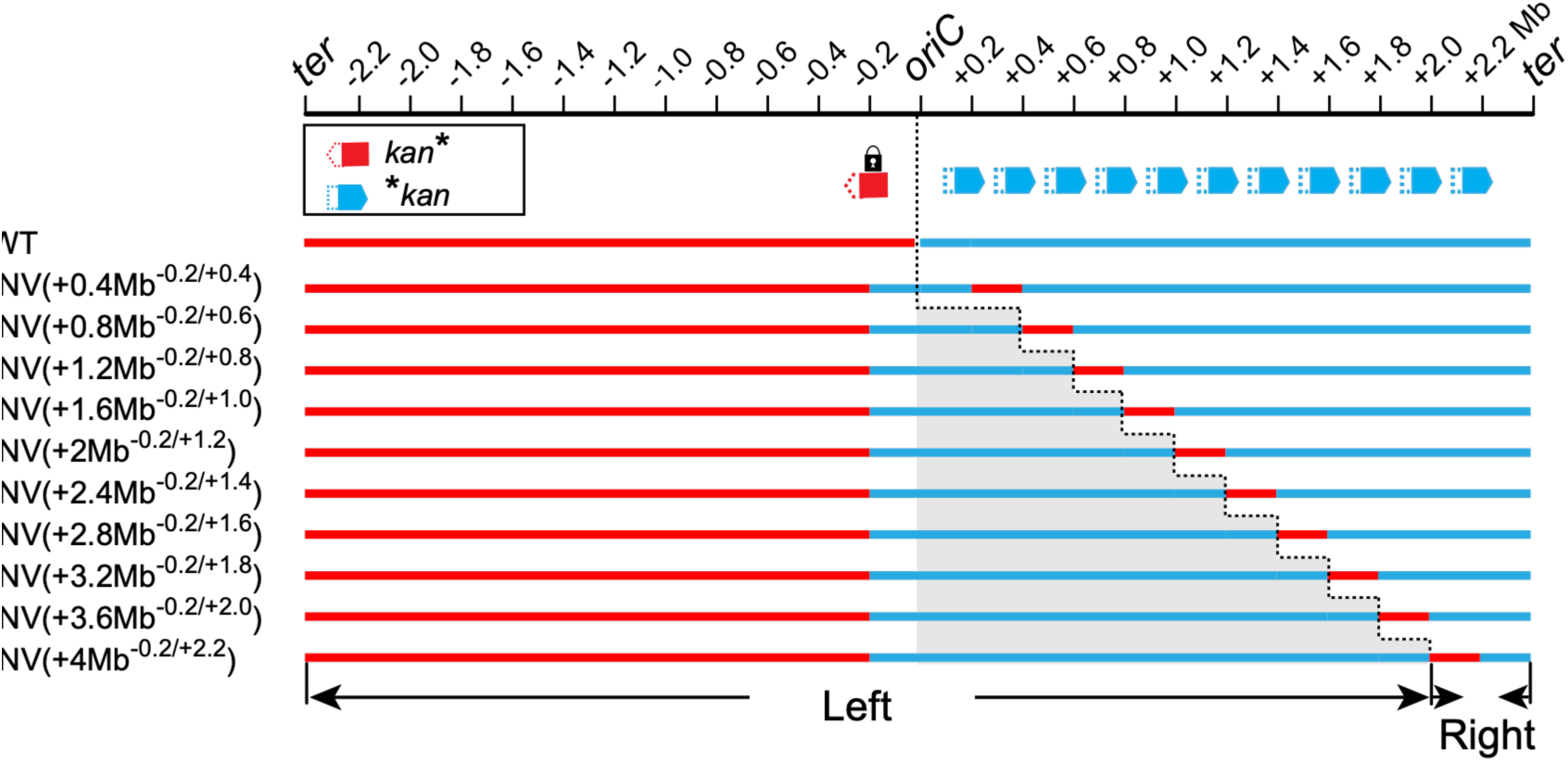
Schematic maps of replichore sizes after chromosomal inversions. Schematic representation of replichore sizes following chromosomal inversion for one of the constructed strain sets. Inversions were generated between a fixed site located at - 0.2 Mb from *oriC* and one of eleven positions on the right replichore, spaced at 0.2 Mb intervals.

**Fig. S2.**
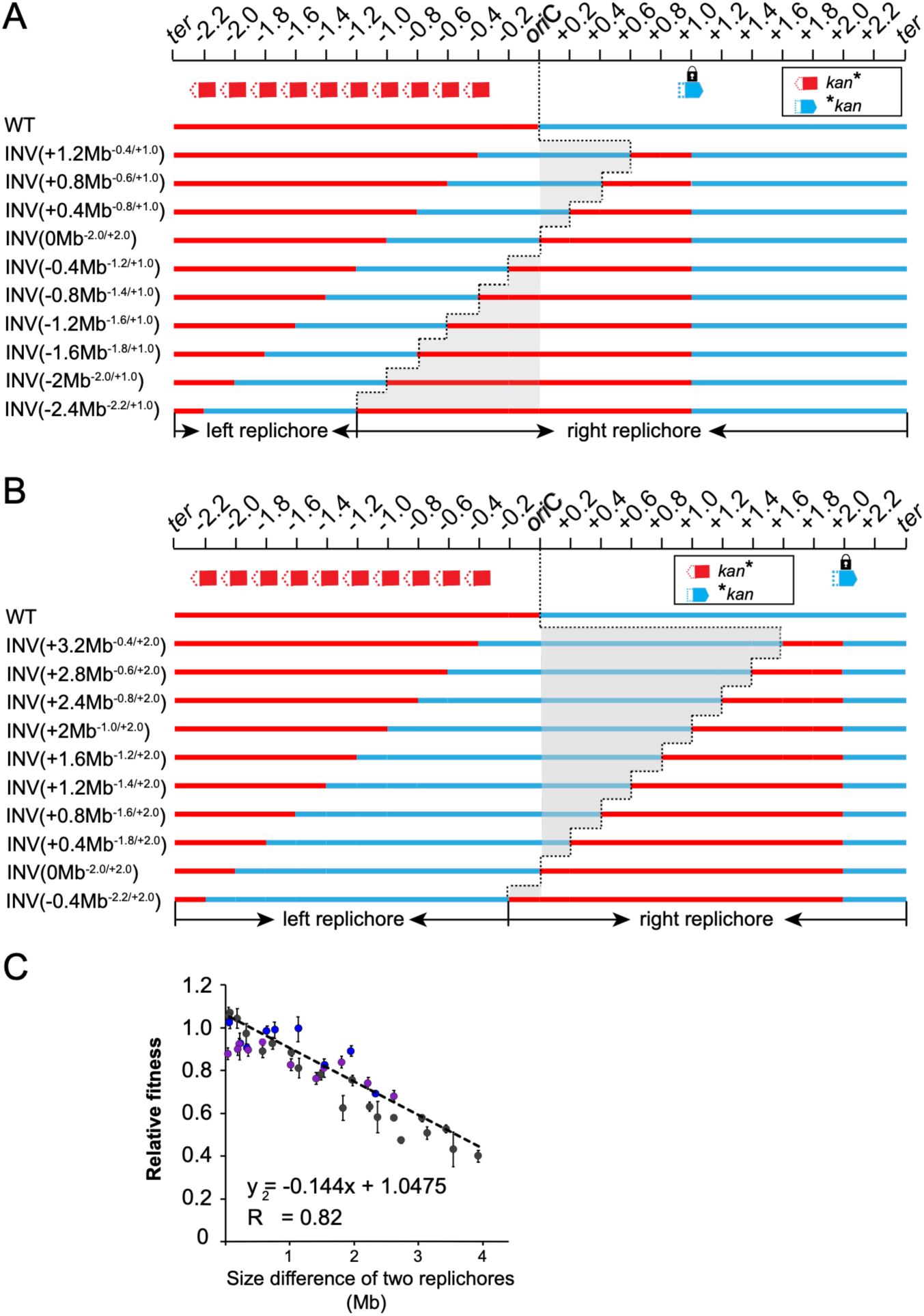
Schematic maps of replichore sizes after chromosomal inversions. (A - B) Schematic representation of replichore sizes following chromosomal inversion. (A) Chromosomal inversions generated betweena fixed site at +1.0 Mb from *ori*C and one of ten positions on the left replichore, spaced 0.2 Mb intervals. (B) Chromosomal inversions generated between a fixed site at +2.0 Mb from *ori*C and one of ten different positions on the left replichore, spaced 0.2 Mb intervals. (C) Correlation between increasing replichore size asymmetry and bacterial fitness. Black dots indicate the relative fitness of inversion mutants at fixed sites at +0.2 Mb and -0.2 Mb; purple dots indicate mutants with fixed sites at +1.0 Mb; blue dots indicate mutants with fixed sites at +2.0 Mb.

**Fig. S3.**
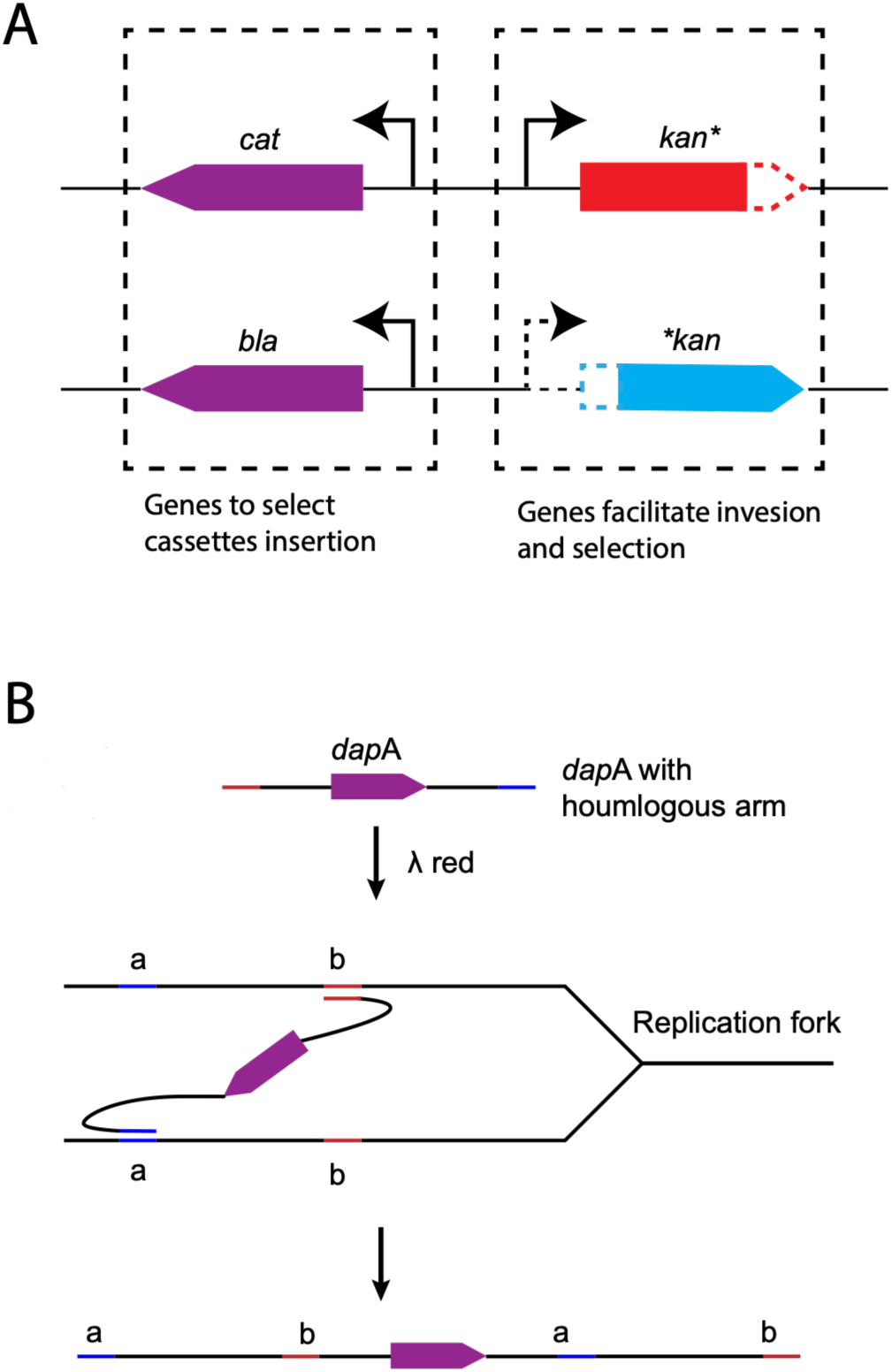
Strain constructions. (A) Design of two cassette sets, *cat*-*kan*\* and *bla-*kan*, enabling both selection of cassette insertion and of chromosomal inversions. The chloramphenicol (*cat*) or ampicillin (*bla*) resistance genes served as selectable markers for cassette insertion at designated chromosomal sites. *kan*\* and \**kan* fragments are truncated versions of the kanamycin (*kan*) resistance gene that retain 645 bp of homologous sequence, which facilitates homologous recombination and chromosomal inversion. (B) Overview of the strategy used to construct chromosomal duplications.

## Tables

**Table S1.**
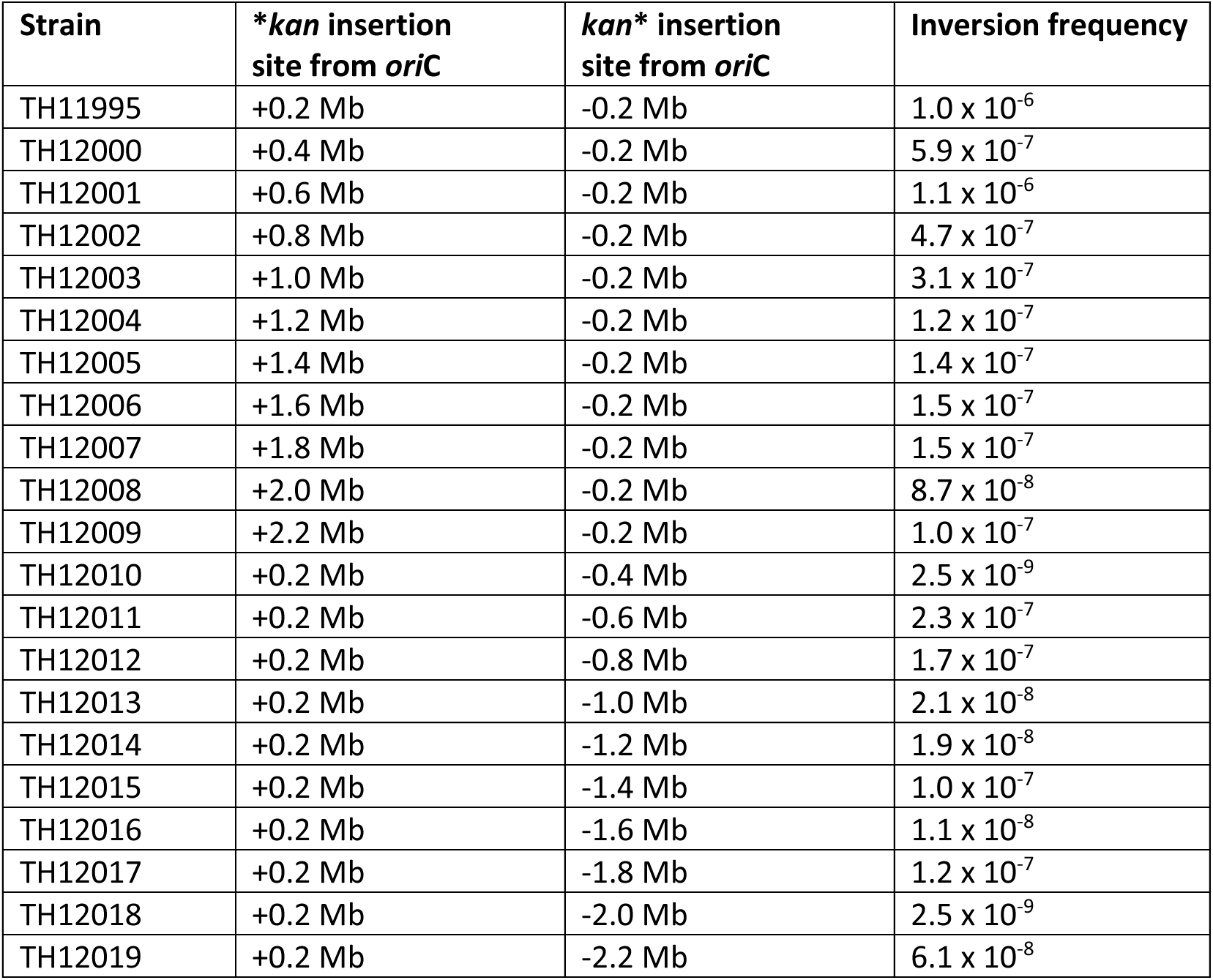
Overview over inversion frequencies.

**Table S2.**
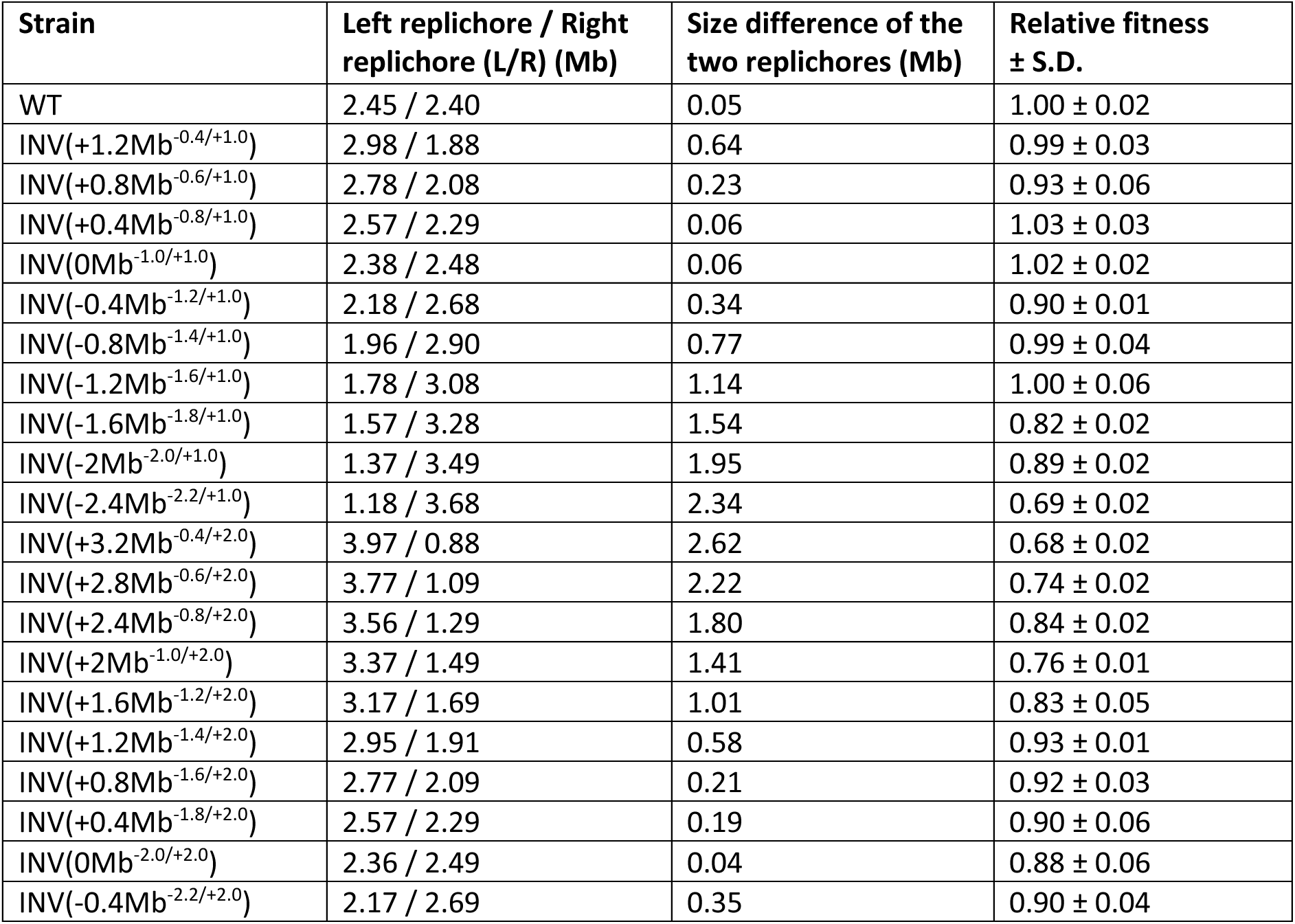
Overview over replichore size differences and fitness effects in inversion strain with fixed insertions site at chromosomal positions +1.0 Mb and +2.0 Mb.

| Strain | Left replichore / Right replichore (L/R) (Mb) | Size difference of the two replichores (Mb) | Relative fitness ± S.D. |
| --- | --- | --- | --- |
| WT | 2.45 / 2.40 | 0.05 | 1.00 ± 0.02 |
| INV(+1.2Mb<sup>-0.4/+1.0</sup>) | 2.98 / 1.88 | 0.64 | 0.99 ± 0.03 |
| INV(+0.8Mb<sup>-0.6/+1.0</sup>) | 2.78 / 2.08 | 0.23 | 0.93 ± 0.06 |
| INV(+0.4Mb<sup>-0.8/+1.0</sup>) | 2.57 / 2.29 | 0.06 | 1.03 ± 0.03 |
| INV(0Mb<sup>-1.0/+1.0</sup>) | 2.38 / 2.48 | 0.06 | 1.02 ± 0.02 |
| INV(-0.4Mb<sup>-1.2/+1.0</sup>) | 2.18 / 2.68 | 0.34 | 0.90 ± 0.01 |
| INV(-0.8Mb<sup>-1.4/+1.0</sup>) | 1.96 / 2.90 | 0.77 | 0.99 ± 0.04 |
| INV(-1.2Mb<sup>-1.6/+1.0</sup>) | 1.78 / 3.08 | 1.14 | 1.00 ± 0.06 |
| INV(-1.6Mb<sup>-1.8/+1.0</sup>) | 1.57 / 3.28 | 1.54 | 0.82 ± 0.02 |
| INV(-2Mb<sup>-2.0/+1.0</sup>) | 1.37 / 3.49 | 1.95 | 0.89 ± 0.02 |
| INV(-2.4Mb<sup>-2.2/+1.0</sup>) | 1.18 / 3.68 | 2.34 | 0.69 ± 0.02 |
| INV(+3.2Mb<sup>-0.4/+2.0</sup>) | 3.97 / 0.88 | 2.62 | 0.68 ± 0.02 |
| INV(+2.8Mb<sup>-0.6/+2.0</sup>) | 3.77 / 1.09 | 2.22 | 0.74 ± 0.02 |
| INV(+2.4Mb<sup>-0.8/+2.0</sup>) | 3.56 / 1.29 | 1.80 | 0.84 ± 0.02 |
| INV(+2Mb<sup>-1.0/+2.0</sup>) | 3.37 / 1.49 | 1.41 | 0.76 ± 0.01 |
| INV(+1.6Mb<sup>-1.2/+2.0</sup>) | 3.17 / 1.69 | 1.01 | 0.83 ± 0.05 |
| INV(+1.2Mb<sup>-1.4/+2.0</sup>) | 2.95 / 1.91 | 0.58 | 0.93 ± 0.01 |
| INV(+0.8Mb<sup>-1.6/+2.0</sup>) | 2.77 / 2.09 | 0.21 | 0.92 ± 0.03 |
| INV(+0.4Mb<sup>-1.8/+2.0</sup>) | 2.57 / 2.29 | 0.19 | 0.90 ± 0.06 |
| INV(0Mb<sup>-2.0/+2.0</sup>) | 2.36 / 2.49 | 0.04 | 0.88 ± 0.06 |
| INV(-0.4Mb<sup>-2.2/+2.0</sup>) | 2.17 / 2.69 | 0.35 | 0.90 ± 0.04 |

**Table S3.**
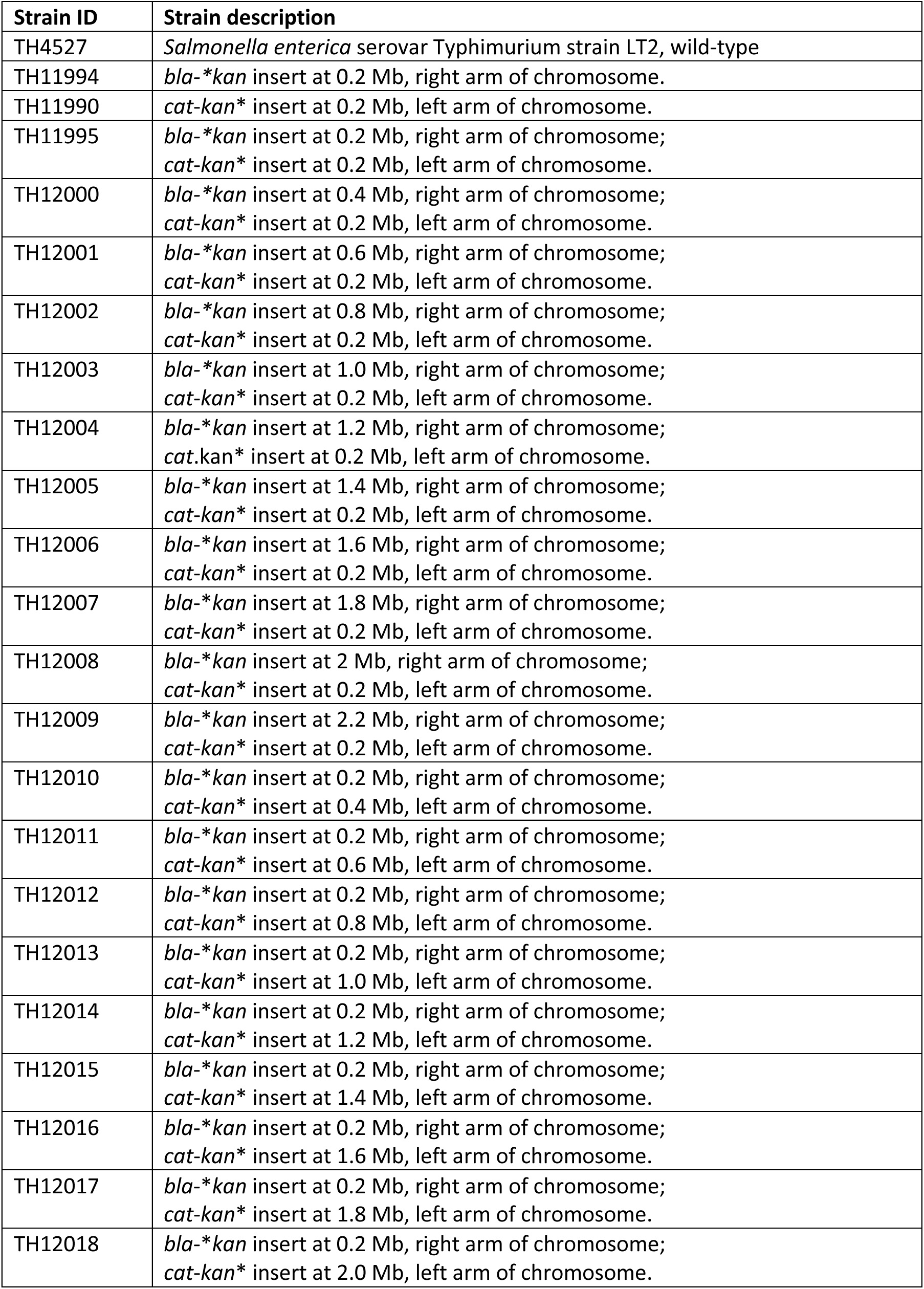
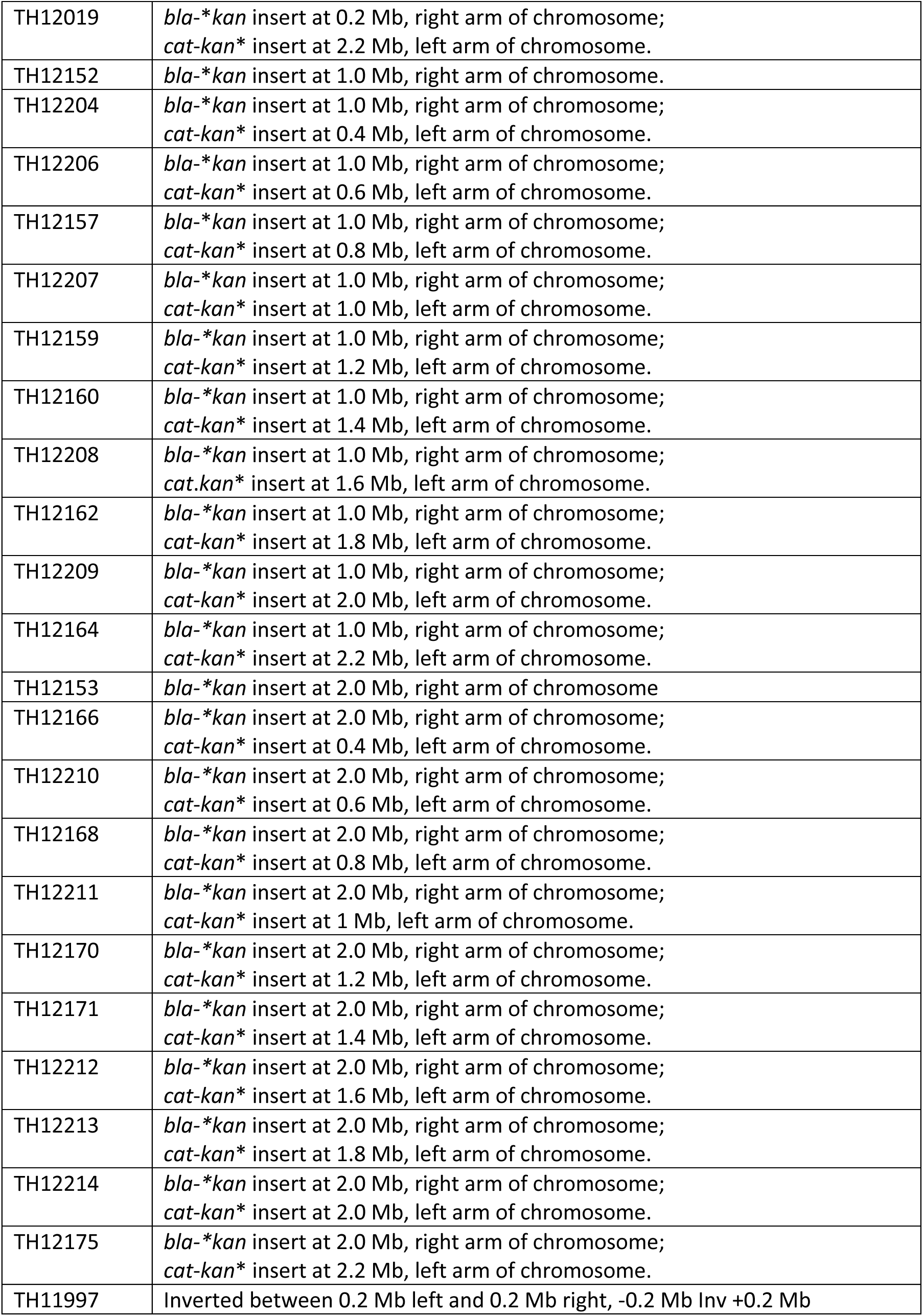
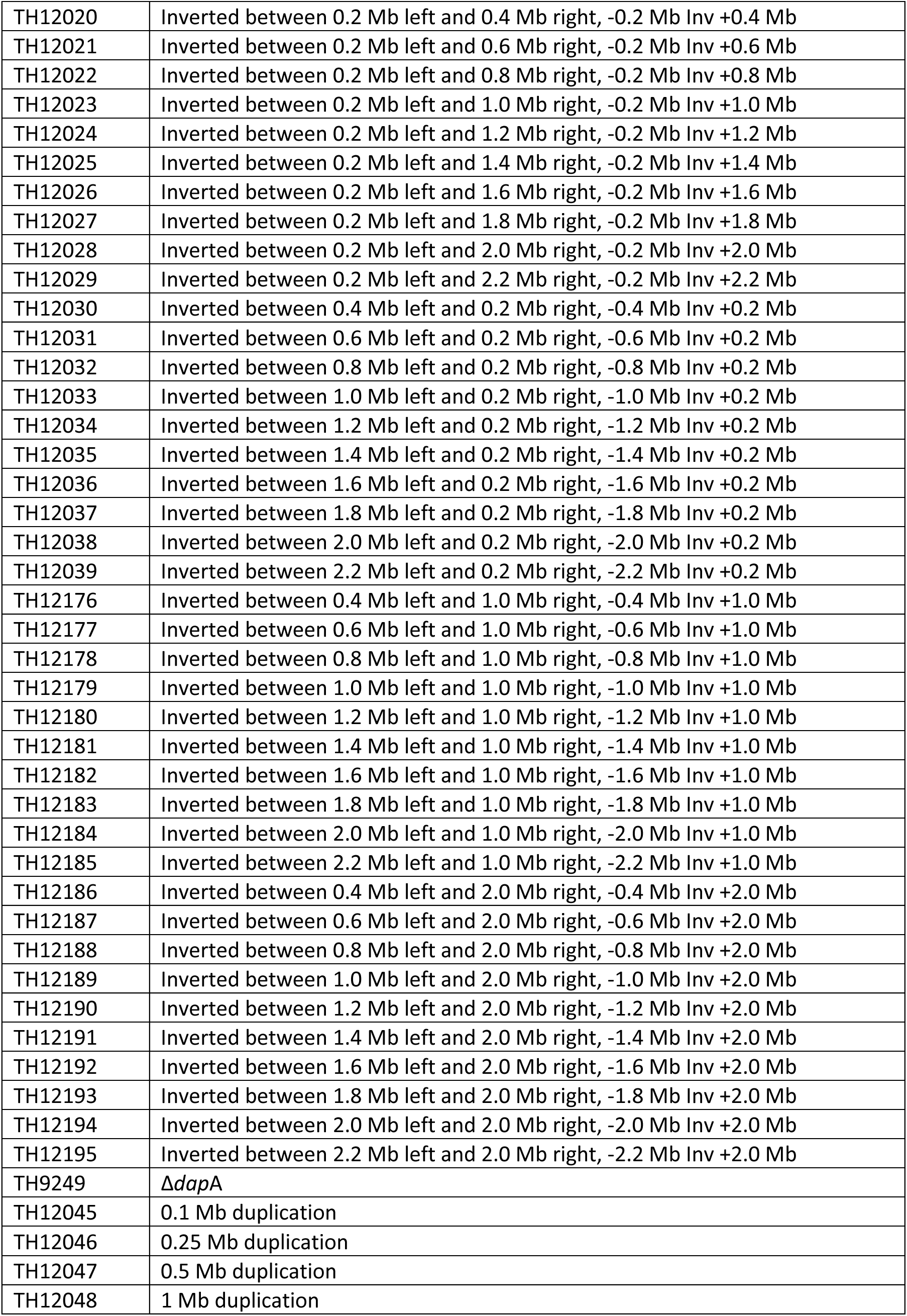
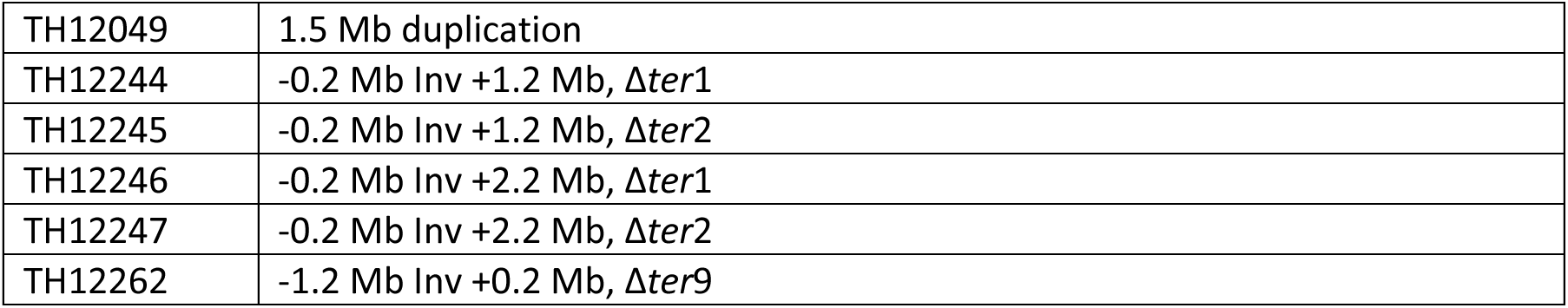
Strain list.

**Table S4.**
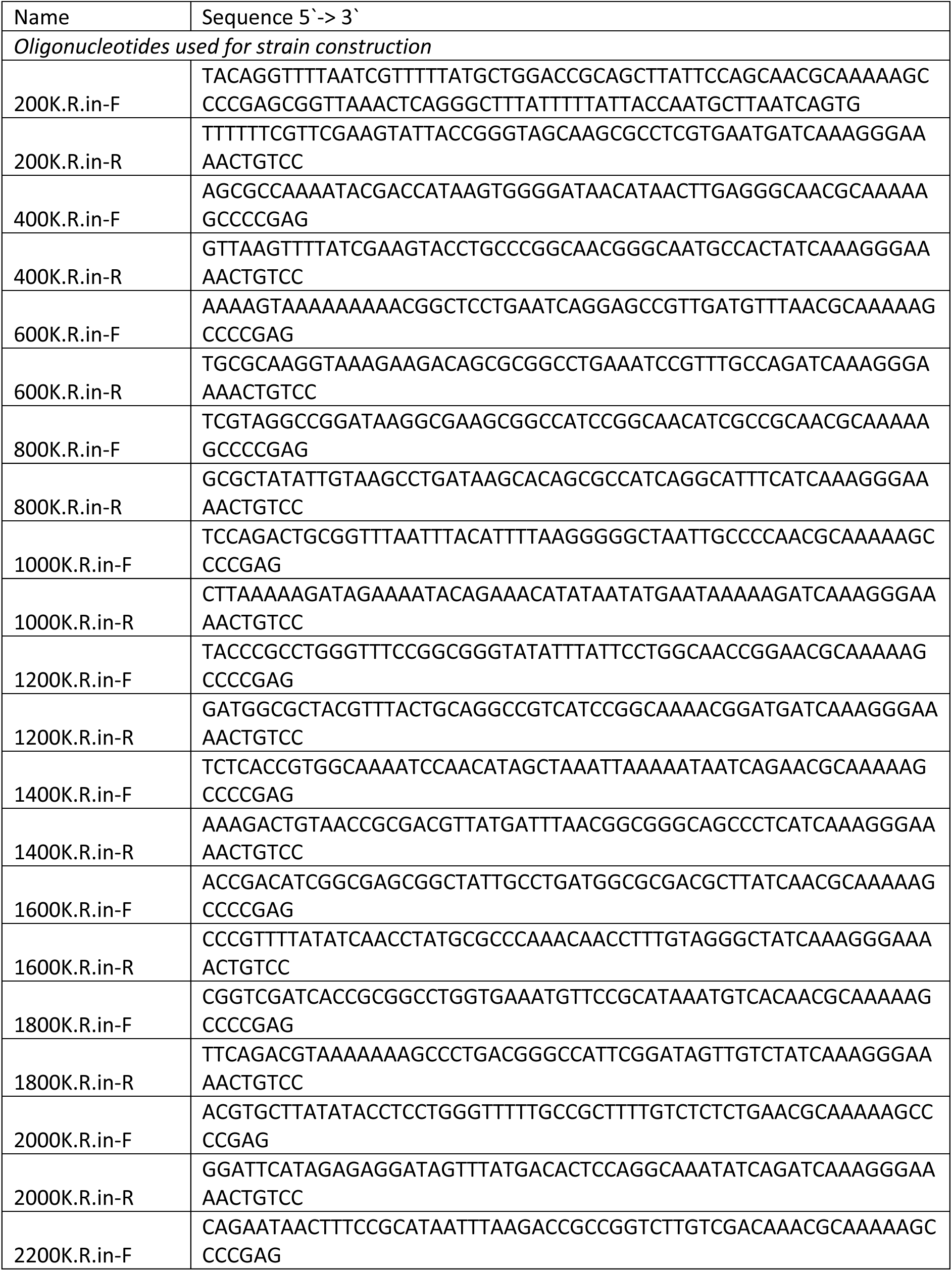
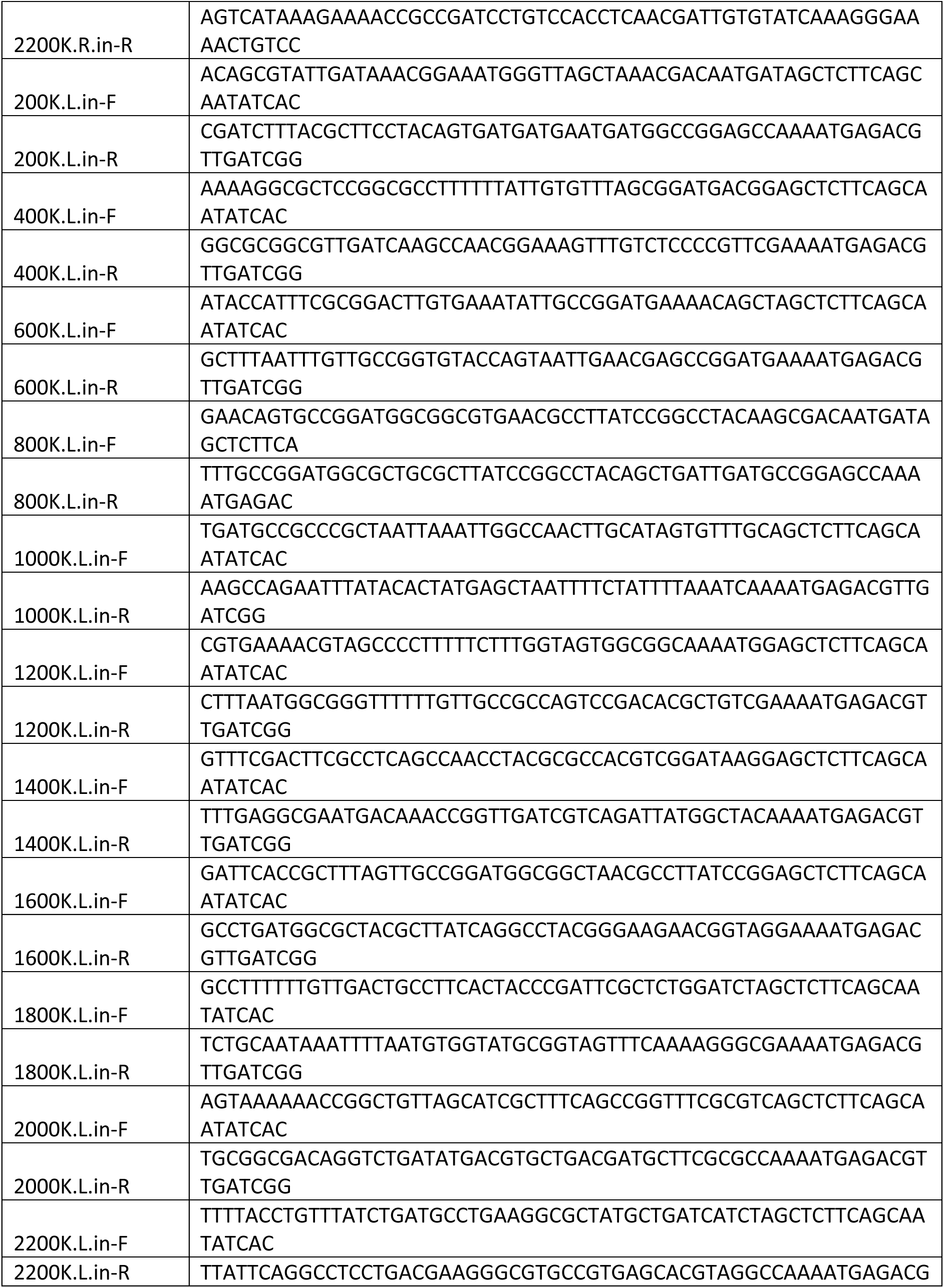
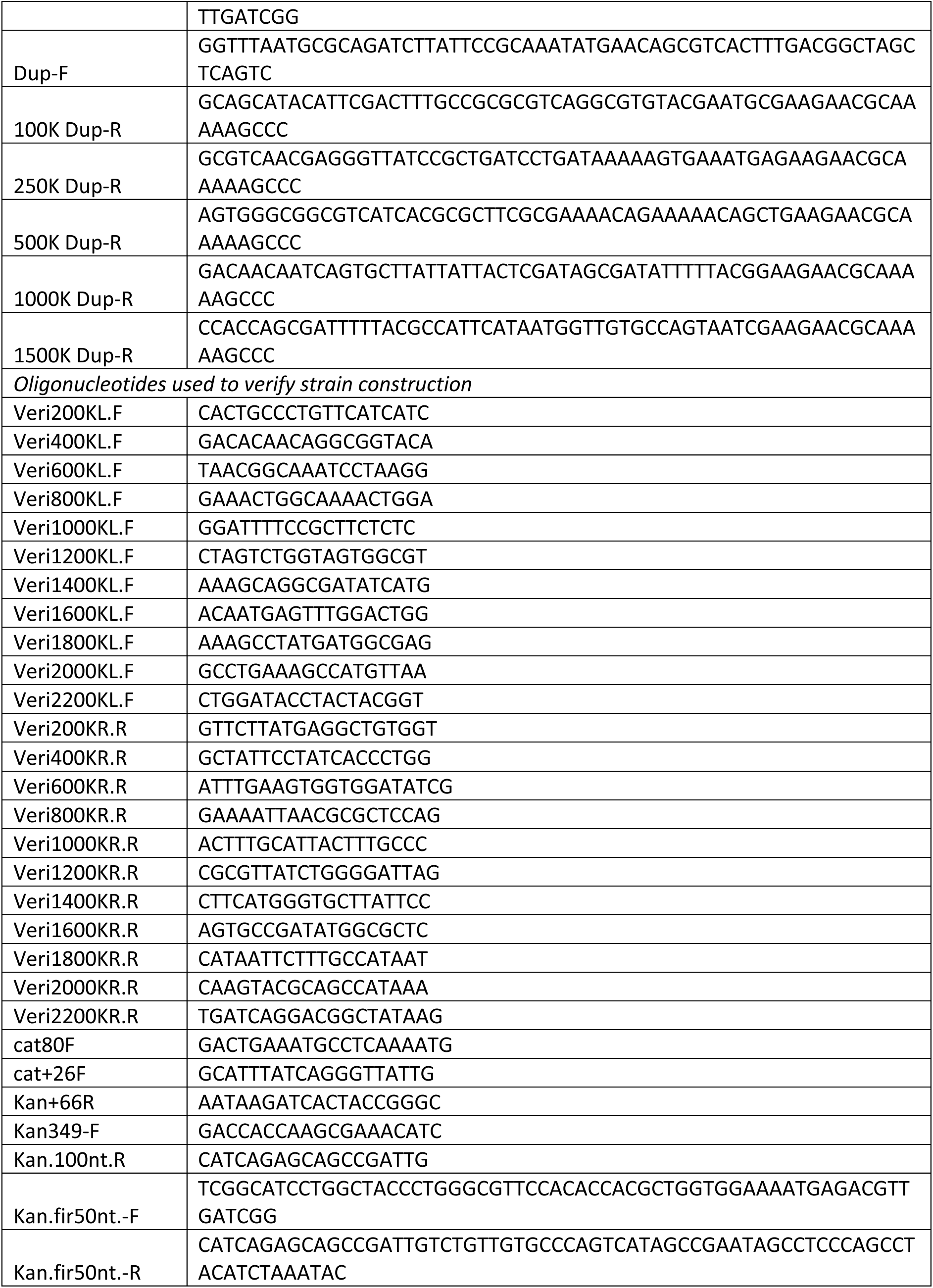
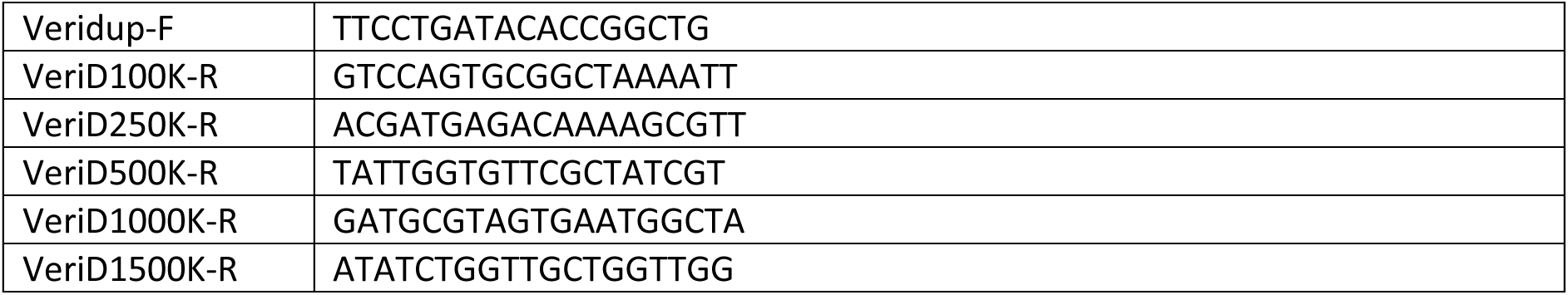
List of oligonucleotides.

## Notes

### Competing Interest Statement

The authors have declared no competing interest.

